# An approach to characterize mechanisms of action of anti-amyloidogenic compounds *in vitro* and *in situ*

**DOI:** 10.1101/2023.12.18.572111

**Authors:** P. Stalder, T. Serdiuk, D. Ghosh, Y. Fleischmann, L. Malinovska, A. Davranche, W. Haenseler, C. Boudou, E. Tsika, A. Ouared, J. Stöhr, R. Melki, R. Riek, N. de Souza, P. Picotti

## Abstract

Aggregation of amyloidogenic proteins is associated with neurodegenerative disease and its modulation is a focus of drug development efforts. However, the physicochemical properties and structural heterogeneity of amyloidogenic proteins hinder the mechanistic understanding of anti-amyloidogenic compounds. Further, modes of interaction with amyloidogenic proteins are often probed *in vitro* using purified protein samples, even though these models may not capture *in vivo* protein structures and do not enable identification of off-target effects. We have developed a modular structural proteomic pipeline based on limited proteolysis coupled to mass spectrometry (LiP-MS) with improved, amino acid level-resolution, to probe the mechanism of action of anti-amyloidogenic compounds. We demonstrate our approach by analysing the interactions of six known or putative anti-amyloidogenic compounds and the amyloid binder Thioflavin T (ThT) with different structural forms of the amyloidogenic Parkinson’s disease (PD) protein α-Synuclein. Our approach enables determination of putative interaction sites, identification of whether interactions are covalent or non-covalent, and crucially, can probe for interactions of compounds with physiological structures of α-Synuclein in complex cell and tissue extracts and identify off-targets. *In vitro* analyses with our pipeline showed that the green tea polyphenol EGCG induces an N- and C-terminus-dependent compaction of the unstructured α-Synuclein monomer, detected preferential interactions of ThT with the N-terminus of α-Synuclein fibrils compared to the amyloid core, and showed that the most potent inhibitors of aggregation in our study (EGCG, baicalein and AC Immune compound #2) induced similar non-fibrillar end structures despite different interactions with α-Synuclein monomers. Importantly, in mammalian cell lysates, α-Synuclein was either a low-affinity target (for EGCG and Baicalein) or did not show evidence of compound interaction (for ThT and doxycycline) in our experimental conditions, despite both monomeric and fibrillar forms interacting with these compounds *in vitro*. For EGCG, we validated this result in postmortem brain homogenates from PD patients. These *in situ* analyses identified many additional putative cellular targets of Doxycycline, EGCG, Baicalein and ThT, suggesting that their effects in cellular or animal models of neurodegeneration are likely due to interactions with proteins other than α-Synuclein and showing that anti-amyloidogenic compounds should be analyzed *in situ* as well as *in vitro*. Our modular pipeline will enable *in situ* screening of drugs and PET tracers for amyloid aggregates of interest as well as detailed mechanistic studies of compound action *in vitro*.

## Introduction

Amyloidogenic proteins can undergo a structural transition to form highly ordered, cross β-sheet rich amyloid fibrils. Although amyloid aggregation can be functional and conserved^1^, numerous amyloidogenic proteins are implicated in disease. In particular, neurodegenerative diseases such as Alzheimer’s (AD) and Parkinson’s disease (PD) are believed to involve the aggregation of specific amyloidogenic proteins, which have been found in inclusions in post-mortem brains of patients and genetically linked to these diseases^2–5^. Preventing aggregation or eliminating aggregates of disease relevant proteins is therefore a strong focus of drug development efforts, but with limited success so far.

Screening for anti-amyloidogenic compounds is typically done either by probing *in vitro* for modification of the aggregation process of a given protein^6–8^, or by screening for reversal or modification of phenotypes in cellular models of the disease^9^ (e.g., formation of foci containing aggregated proteins). However, in both screening approaches, there are limitations in deciphering the mechanism of action of compounds that show an effect. Specifically, a compound could have its effect via covalent or noncovalent binding or otherwise interfering with the monomeric, amyloid fibrillar, or other structural states of the amyloidogenic protein. Compound mechanism of action, in particular knowing which structural state is targeted, is key information for design of validation and selectivity studies as well as for structure-activity relationship (SAR)-based modifications of a drug to increase potency and reduce side effects^10,11^. Further, fibril structures prepared *in vitro* may not recapitulate structures that are formed in patients^12–17^, and effects in cells could be indirect and due to binding to other targets. There is therefore a strong need for approaches to characterize anti-amyloidogenic drugs that can reveal which structural state of an amyloidogenic protein is bound by the drug and what the binding sites are; whether and how the compound affects structural transitions of amyloidogenic proteins over time; whether the drug interacts with the aggregation prone protein in patient samples; and whether it targets other cellular proteins that could have indirect or unwanted effects.

Aggregation-prone proteins possess intrinsic physicochemical properties that make mechanistic studies of compound action very challenging. The non-aggregated, monomeric form of amyloidogenic proteins is often highly dynamic and unstructured^18–21^, so that structural analysis and deconvolution of compound binding are more complex than for proteins with well-defined 3D structures. Once amyloid fibrils are formed, their insolubility and size interfere with a variety of techniques. Recent advances in cryo-electron microscopy and solid-state NMR have yielded fibril structures, but analysis of drug binding as performed for example for the protein Tau and the compound Epigallocatechin gallate (EGCG)^22^ is time intensive, and as yet few such studies have been reported^23–25^. Further, most structural techniques are limited to purified proteins outside of their native environment or require electroporation of purified proteins^26^ and are therefore limited to the analysis of one protein at a time. A robust technique to characterize anti-amyloidogenic drug mechanism of action that can be applied both in purified and in complex settings, in a structurally aware manner, is still missing.

Here, we show that LiP-MS can be used to characterize the mechanism of action of anti-amyloidogenic compounds *in vitro* and directly in cell and tissue extracts. The approach employs a modular experimental pipeline, relying on a new high-resolution version of LiP-MS, a structural proteomic approach that probes protein structure based on preferential protease accessibility of flexible and surface exposed regions of proteins^27^. Cleavage products are measured by mass spectrometry, yielding information about altered surface accessibility of proteins upon a perturbation such as anti-amyloidogenic compound treatment. Our high-resolution LiP-MS approach enables the extraction of interaction sites and structural information at near-amino acid resolution. Further, it allows assessment of whether a compound of interest interacts with a specific structural state *in vitro* (e.g., monomeric or fibrillar states of an amyloidogenic protein) and of how the compound affects the structural evolution of the protein over time. The approach can provide proteome-level data from cell and tissue extracts, thus enabling assessment of compound interactions with the endogenous forms of the amyloidogenic protein but also with other potential protein targets.

We applied this pipeline to characterize the mode of action of a set of anti-amyloidogenic compounds (EGCG, Baicalein, doxycycline, Fasudil, and two proprietary anti-amyloidogenic compounds from drug discovery efforts) and of the amyloid binder ThT, using the PD-associated protein α-Synuclein as a test case. Applied to an aggregation time course *in vitro*, our approach revealed that the most potent inhibitors of aggregation (EGCG, Baicalein, ACI compound # 2) resulted in structures that were distinct from both monomers and fibrils and may also represent a mixture of states. We identified interactions with monomer and/or fibrillar forms of α-Synuclein *in vitro* for most of the compounds; for instance, EGCG induced a compaction of α-Synuclein monomer via interactions with the N- and C-terminus of the protein but caused structural changes in α-Synuclein fibrils at the N-terminus and the C-terminal region of the aggregation core. Surprisingly, Fasudil had no structural effects on either form of α-Synuclein and ThT interacted with the N-terminus of α- Synuclein fibrils but only weakly with the aggregation core i.e., the non-amyloid component (NAC).

Applied in the context of a mammalian cell lysate, our approach showed only low-affinity interaction, or no interaction at all, of EGCG, Baicalein, doxycycline and ThT with overexpressed α-Synuclein, while concomitantly identifying numerous other putative cellular interactors of these compounds, consistent with prior classification of EGCG and baicalein as pan-assay interference (PAIN) compounds^28^. Our data indicate that effects of these compounds in *in situ* models of neurodegeneration must be due to interactions with proteins other than α-Synuclein, and argue strongly that screening and characterization of anti-amyloidogenic compounds and PET tracers should be carried out *in situ*.

## Results

### An amino acid-resolution analytical approach for high-coverage LiP-MS data

We first asked if we could use LiP-MS to derive structural information about anti-amyloidogenic or amyloid-binding compound effects on purified proteins *in vitro*. LiP-MS analysis of purified proteins typically results in the detection of a large number of partially overlapping peptides along a protein sequence. To take advantage of the resulting high-coverage data and increase the structural resolution of LiP-MS, we developed a new analytical approach. As a test case, we compared the LiP fingerprints of intrinsically disordered α-Synuclein monomers and α-Synuclein amyloid fibrils *in vitro*. We first analyzed the data with the classical LiP-MS pipeline which achieves peptide-level resolution. As expected, differential analysis showed that many peptides were significantly different in abundance (|log_2_FC| >1, q-value < 0.05) between the two structures (**Figure 1A**). When significantly changing peptides were mapped along the linear sequence of α-Synuclein, almost the entire sequence was conformationally different (**Figure 1C**), because of the large structural changes between the two forms of the protein. In such a case, a peptide-level analysis gives limited information about detailed structural changes across different regions of the protein. We therefore designed an alternative analysis strategy to combine information from multiple peptides mapping to the same region and gain higher-resolution insight into the conformational landscape of α-Synuclein.

**Figure 1.**
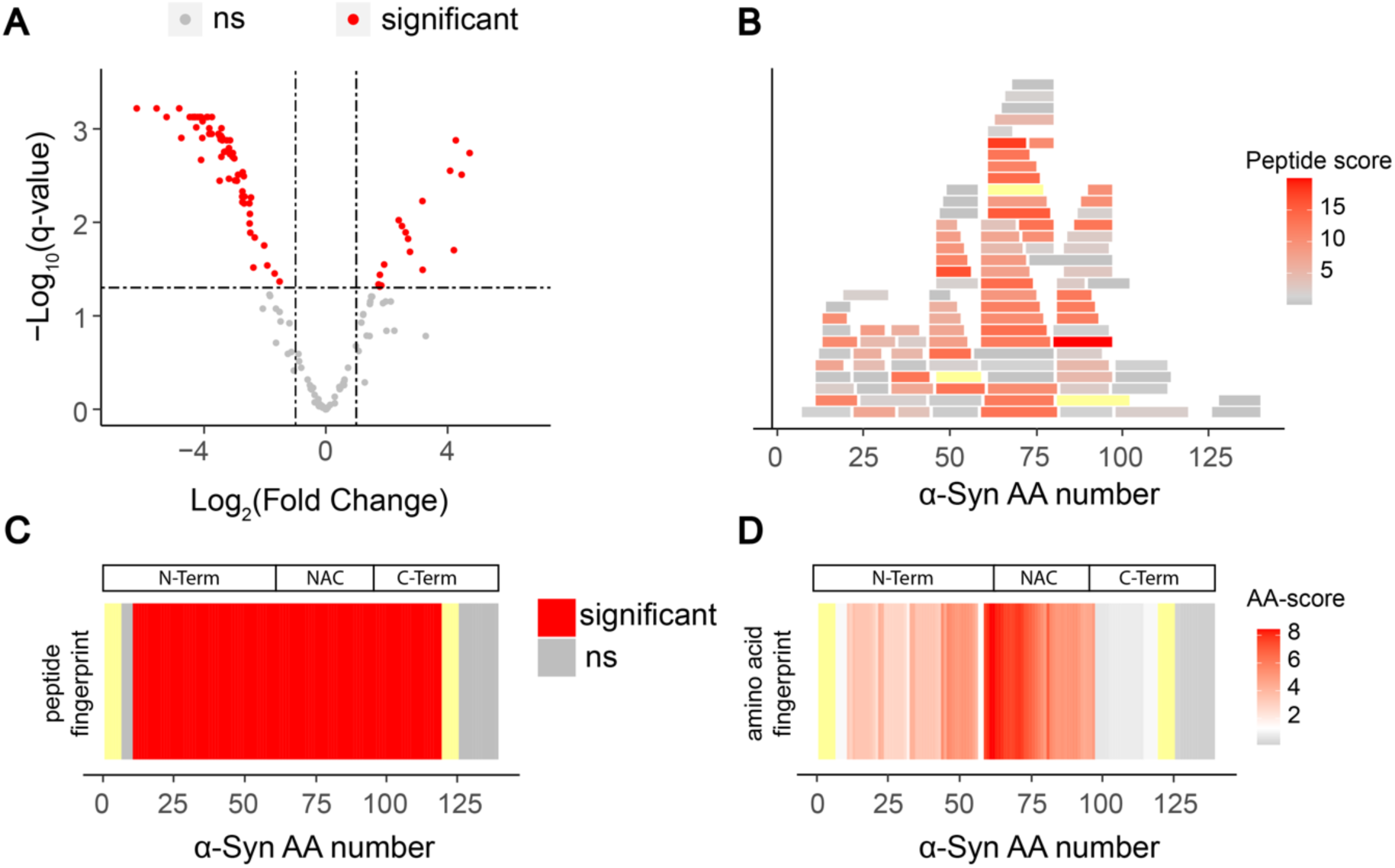
A new amino acid-centric analysis approach for higher-resolution structural comparison with LiP-MS. **A** Volcano plot comparing LiP peptide abundances generated upon limited proteolysis of α-Synuclein monomers and fibrils. **B** All detected and quantified LiP peptides and their corresponding scores (-log_10_(q-value) x |log_2_(fold change)|) were mapped along the α-Synuclein sequence. The scale reflects the score for every peptide. The more intense the red colour, the higher the score. For all panels, not significant is shown in grey, not detected in yellow. **C** Structural fingerprint comparing α-Synuclein monomer and fibril after applying the classical LiP-MS data analysis pipeline. Significant regions (|log_2_FC| >1, q-value < 0.05) in red. **D** Structural fingerprint comparing α-Synuclein monomer and fibril upon scoring changes per amino acid. The scale indicates the score per amino acid. The significance threshold of −log_10_(0.05) x |log_2_(2)| is shown in white, with red indicating higher scores. The more intense the red colour, the higher the score. N-term, N-terminus; NAC, non-amyloid β component (aa 61-95); C-term, C-terminus.

Similar to approaches used in RNA sequencing analysis^29^, we first assigned a score to every peptide, corresponding to the|log_2_(fold change)| in peptide intensity between the two conditions times the log10(q-value) as a measure of statistical significance. The larger and more significant the fold change, the higher the score for a given peptide. Next, we overlapped all the peptides (**Figure 1B**) and calculated the mean value of this score per amino acid position. Aggregating the data per amino acid across the entire protein sequence then yielded a detailed structural fingerprint of α-Synuclein monomers compared to fibrils (**Figure 1D**). At a comparable significance threshold as the peptide level analysis (|log_2_(fold change)|> 1; q-value < 0.05), we now observed that the NAC core region^30^ until amino acid 95 showed strong structural differences between the two forms of the protein, as expected since this region is arranged in cross β-sheets in the fibril but not the monomeric form^31^. Also as expected, the C-terminus, which is known to be flexible in both structural states of the protein, did not differ between monomeric and fibrillar α-Synuclein. Smaller but significant differences were observed for the N-terminus, suggesting that a fraction of the N-termini of α- Synuclein fibrils are structured or have reduced protease accessibility under the buffer conditions used^32^. These results illustrate that our new amino acid centric approach yields the expected patterns for a structural comparison of α-Synuclein monomer and fibril and allows a more fine-grained picture of the structural differences between the two states of the protein.

We further tested whether this amino acid centric analysis could pinpoint small molecule binding sites from high coverage LiP-MS data with improved resolution. To this end, we used LiP-MS data from our previous study^33^ (**Figure 2**), which showed that fructose-1,6-bisphosphate (FBP) binds at the phosphoenolpyruvate (PEP) binding site of Enzyme I of the PEP-dependent sugar phosphotransferase system (ptsI), thereby inhibiting enzyme activity by competitive inhibition. To test the resolution of the amino acid centric approach, we analysed the LiP-MS data comparing the FBP-bound and unbound states of ptsI either with our previous peptide-level analysis or the new amino acid centric approach. We found that our new approach closely mapped the significantly changing amino acids to the known PEP binding cleft of ptsI (**Figure 2C, E**). Our previous peptide-level quantification also allowed mapping of the binding site, but with lower precision (**Figure 2B, D**). We achieved similar results upon analysis of *in silico* LiP-MS data, showing in addition that the amino-acid approach may be more resistant to false positives (**Supplementary Figure 1**). Finally, we used our approach to assess binding of Vitamin D binding protein (GC) to Vitamin D (**Supplementary Figure 2**), Fructose-bisphosphate aldolase A (ALDOA) to fructose bisphosphate (**Supplementary Figure 3**), and Thyroxine Binding Globulin (TBG) to thyroxine (**Supplementary Figure 4**), in all cases re-analysing data from a previous study in which we added the compounds to purified proteins and comparing proteolytic patterns in the presence versus absence of the compound^34^. For ALDOA and GC, the amino acid centric approach improved the resolution of binding site mapping over our previous peptide-centric analysis. In the case of TBG, both versions of our LiP-MS analysis did not detect a structural change in the known thyroxine binding site, due at least in part to limited coverage of this region. Nevertheless, the amino acid scoring approach detected at much higher resolution the known conformational change of the reactive loop of TBG upon thyroxine binding^35^.

**Figure 2.**
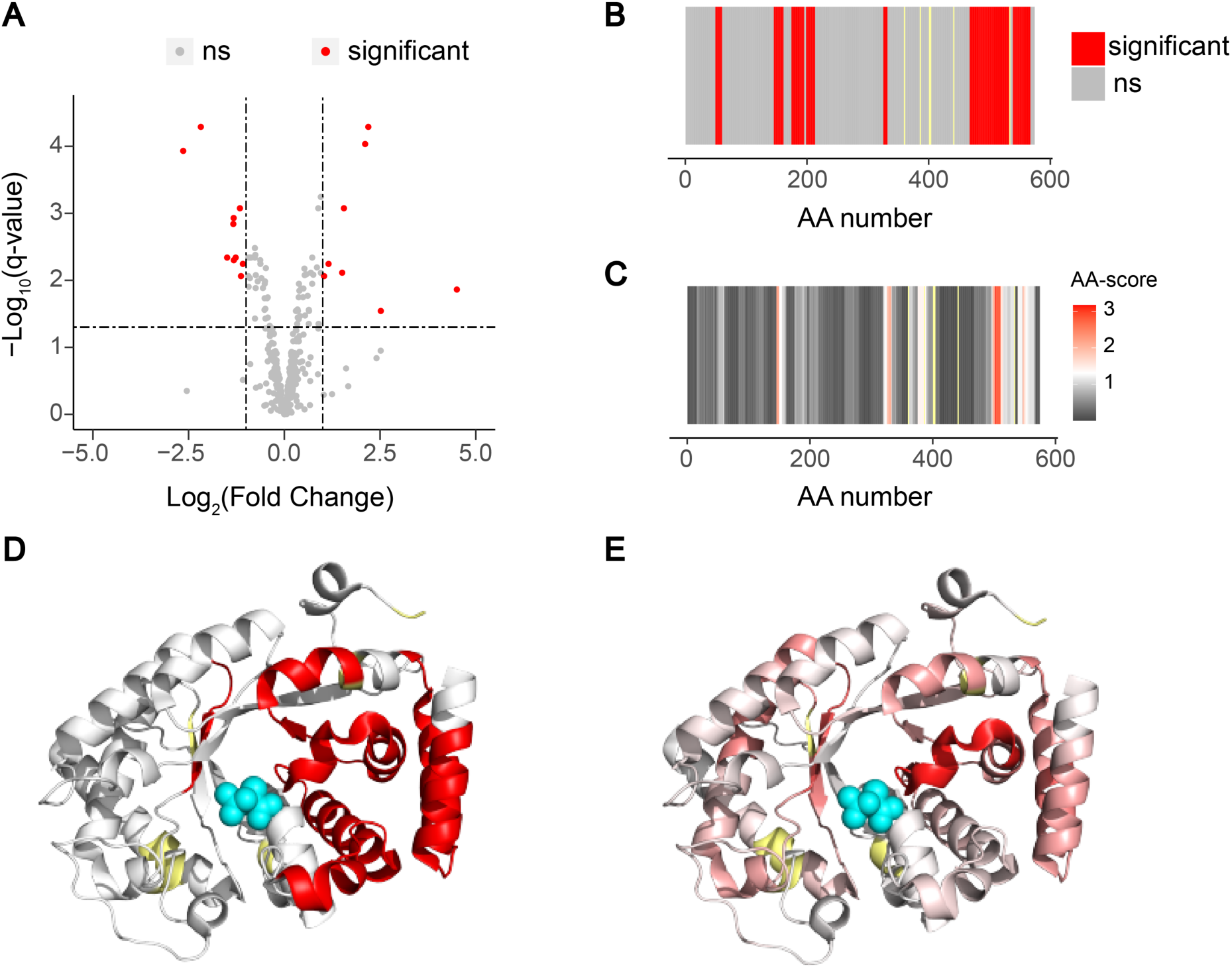
The amino acid centric analysis improves identification of small molecule binding regions. **A** Volcano plot comparing abundances of peptides generated in FBP bound and unbound ptsI. Significant (|log_2_FC| > 1, q-value < 0.05) in red, not significant in grey. **B** Fingerprint of the classical LiP-MS data analysis pipeline. Significant regions in red, not significant in grey, not detected in yellow. **C** Fingerprint upon scoring changes per amino acid. The scale indicates the score per amino acid. The significance threshold of −log_10_(0.05) x |log_2_(2)| is shown in white, with red indicating higher scores. The more intense the red colour, the higher the score. Not significant in grey. Not detected in yellow. **D** Peptide centric fingerprint mapped on the ptsI structure (PDB: 2xz7). PEP in cyan. **E** Amino acid centric fingerprint mapped on the ptsI structure (PDB: 2xz7). PEP in cyan.

High-resolution amino acid-centric analysis of LiP-MS data requires high sequence coverage, which is typically achieved *in vitro* on purified proteins, but not for all proteins in complex backgrounds. Nevertheless, we also tested the amino acid-centric analysis on a dataset in which a yeast lysate was treated with the well-studied small molecule rapamycin^36^. Compared to the *in vitro* data discussed earlier (i.e., α-Synuclein, ptsI, vitamin D binding protein, ALDOA, TBG; Figs. 1-2, Supp Figures 2-4), where the median sequence coverage per protein was 95.1% (**Supplementary Figure 5A**), the proteome-wide data from the rapamycin treated yeast lysate showed a median protein sequence coverage of 14.13% (**Supplementary Figure 5B**) and the main target of rapamycin, FPR1, was covered at 69.3% (**Supplementary Figure 5B**). Even though the amino acid-centric analysis did not in this case increase the resolution of binding site mapping, likely due to the insufficient number of detected peptides, it was still more informative than the classical pipeline since regions with higher scores preferentially mapped to the known binding site of rapamycin (**Supplementary Figure 5C-F**). The amino acid centric analysis can therefore also be informative for proteome wide data since it allows ranking regions by the strength of the effect of a small molecule, but protein coverage will strongly influence the performance.

Overall, we have developed an analytical approach for LiP-MS data that much increases the resolution with which we can pinpoint small molecule binding sites and structural alterations in high-coverage data.

### Structural changes of *α*-Synuclein in the presence of anti-amyloidogenic compounds

Employing our high-resolution amino acid centric approach, we analyzed the effects of candidate anti-amyloidogenic compounds on structural changes of α-Synuclein during amyloid fibril formation. We probed the effects of EGCG, Baicalein, Fasudil and Doxycycline as known inhibitors of aggregation, and two proprietary compounds of AC Immune SA (Switzerland) that inhibit α-Synuclein aggregation in vitro and have protective effects on neuronal cultures challenged with α-Synuclein fibrils, here named ACI compound #1 and compound #2. We also included the known amyloid binding compound ThT.

To test the effects of these compounds on α-Synuclein aggregation, we incubated purified α- Synuclein with α-Synuclein seeds in the presence of each of the compounds (140 uM) or a DMSO control. After 17 hours of incubation under constant agitation, fibril formation had reached steady state in the DMSO-only control as measured by ThT emission (**Figure 3A**); we confirmed fibril formation with electron microscopy (**Supplementary Figure 6**). EGCG and Baicalein completely blocked fibril formation as previously reported^37,38^, since there was no increase in ThT intensity throughout the time course. Doxycycline and compound #1 led to a significant reduction of ThT intensity at the end of the time course, but only caused minor changes in the half time of aggregation (**Figure 3B, C**). In contrast, compound #2 inhibited the aggregation but did not show a sigmoidal curve profile, rather yielding a small linear increase in ThT intensity until the end of the experiment. Fasudil did not affect the aggregation, in contrast to published reports^39^ that used an aggregation assay without fibrillar seeds. We note that compound competition with ThT for binding to α- Synuclein cannot be ruled out. Electron microscopy at the end of the time-course revealed the presence of fibrils in all samples (**Supplementary Figure 6A-G**), although these were visually different in compound #2-, EGCG-, Baicalein-treated samples, our analysis was not quantitative, and the latter two samples also contained oligomeric species as previously reported^37,38^. Native PAGE of the samples further indicated that compound #2-, EGCG-, Baicalein-treated α-Synuclein contained remaining monomeric species, comparable to the monomer at time point zero, while most of the monomeric fraction was lost upon treatment with the other compounds (**Supplementary Figure 6H**).

**Figure 3.**
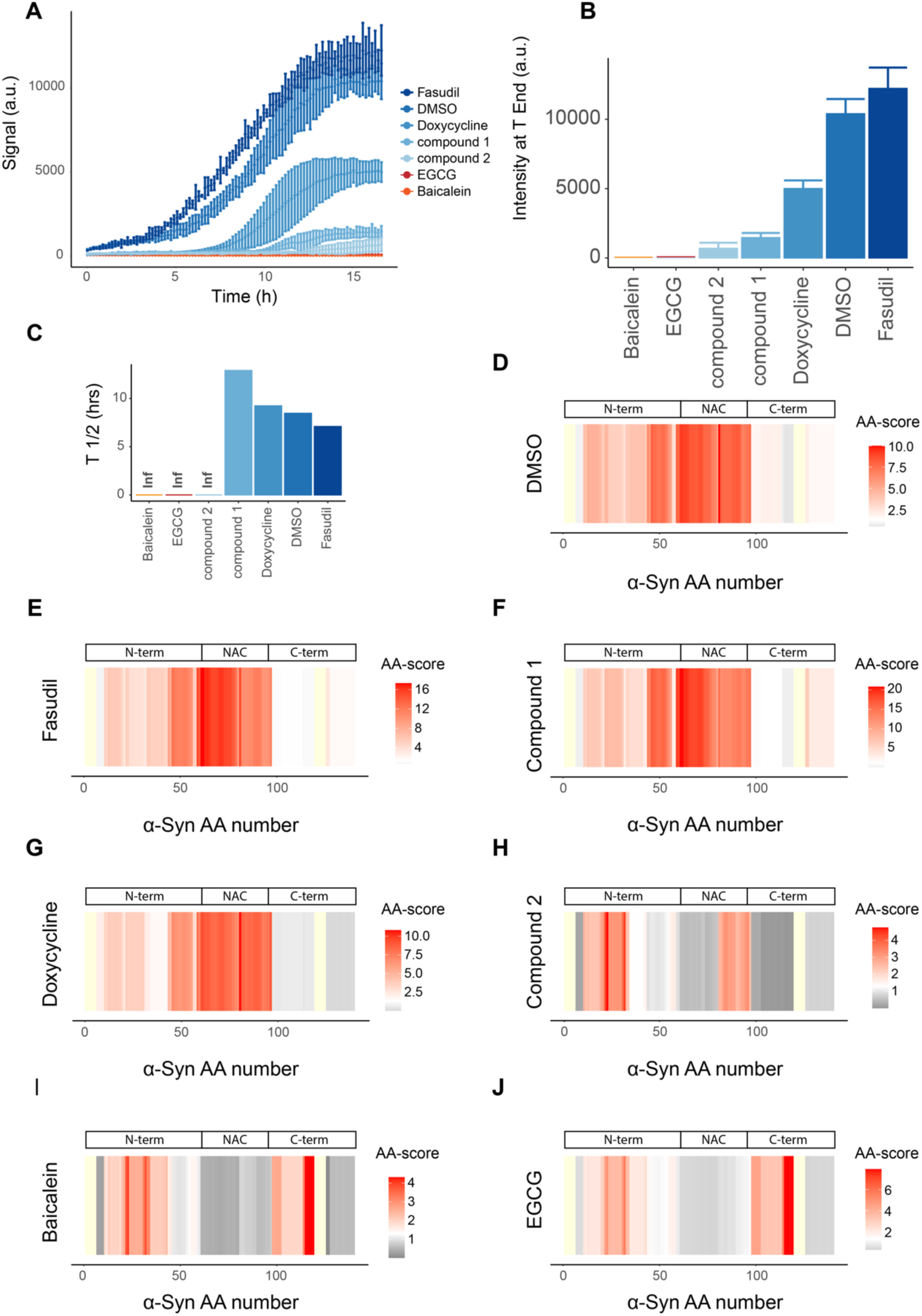
Effects of anti-amyloidogenic compounds on the structure of α-Synuclein upon in vitro aggregation. **A** ThT signal over a 17h α-Synuclein aggregation assay in the indicated conditions (molar ratio compound/α-Synuclein of 5.7). **B** ThT fluorescence signal intensity after 17 hours of incubation (T_End_). **C** Half time (T_½_) of aggregation extracted from ThT fluorescence curve profile. Inf, infinite. **D-J** Structural fingerprints comparing the initial and final α-Synuclein structures under the indicated conditions: DMSO control condition (**D**), Fasudil (**E**), compound #1 (**F**), Doxycycline (**G**), compound #2 (**H**), Baicalein (**I**) and EGCG (**J**). The scale indicates the score per amino acid. The significance threshold of −log_10_(0.05) x log_2_(2) is shown in white, with red indicating higher scores. The more intense the red colour, the higher the score. Not significant in grey. Not detected in yellow. N-term, N-terminus; NAC, non-amyloid β component (aa 61-95); C-term, C-terminus.

To assess in more detail how the structure of α-Synuclein changed over time in the presence of these compounds, we performed a LiP-MS experiment at the start and the end of the time course and derived fingerprints comparing the structures at these two time points for each condition. As expected in the DMSO control sample, the strongest structural changes between the start and end of the time course were in the α-Synuclein aggregation core (aa 61-95)^30^ (**Figure 3D**), which becomes highly structured upon aggregation of α-Synuclein. The structural fingerprint for the Fasudil-treated sample was the same as that of the DMSO control (**Figure 3D, E**), consistent with the fact that Fasudil did not affect the aggregation process in our experiments and indicating that the same fibril structures were formed in the presence of Fasudil and in the control sample. Fibrils formed in the presence of Doxycycline and compound #1 yielded the same structural fingerprint as in the control sample (**Figure 3F, G**), again indicating that similar fibril structures were formed, despite a substantially lower ThT signal in the presence of these compounds; this may suggest interactions of doxycycline or compound 1 with ThT, or competition with ThT for binding to α-Synuclein, or simply a low amount of α-Synuclein fibrils formed under these conditions. We observed a different structural fingerprint for fibrils formed in the presence of compound #2, with less profound changes in the aggregation core relative to the monomeric form, when compared to fibrils formed under control conditions (**Figure 3H**). However, the fingerprint does indicate some changes in the C-terminal part of the aggregation core (aa 81-97)^30^. Most interestingly, structures formed in the presence of EGCG and Baicalein changed in proteolytic accessibility in their very N-and C-terminus, but the aggregation core remained unchanged compared to the monomer (i.e., unstructured) (**Figure 3I, J**). Overall, the most potent inhibitors of aggregation (baicalein, EGCG and compound 2) produced similar structures at the end of the aggregation time course (i.e., structural changes around residue 40 for all compounds, plus residues 90-100 for compound 2, and residues 100-115 for baicalein and EGCG). Since these structural changes relative to monomer largely do not involve the NAC region, these fingerprints indicate that no fibrils were detectable by LiP-MS in EGCG, baicalein and compound #2, consistent with the ThT fluorescence results.

Thus, our approach allows assessment of structures formed by amyloidogenic proteins in the presence of compounds of interest. The aggregation core region of α-Synuclein could be accurately identified by our technique, allowing a direct assessment of the aggregation process.

### Compound interactions with monomeric *α*-Synuclein

To characterize the mechanisms of action of anti-amyloidogenic compounds, it is important to assess whether they interact with specific structural states of the amyloidogenic protein. We thus asked whether the LiP-MS pipeline could detect interaction of our compounds with monomeric α-Synuclein. We prepared monomeric α-Synuclein, verified its purity by blue native PAGE (**Supplementary Figure 7A**), and compared its protease accessibility in the presence and absence of the compounds (1:100 molar ratio) using LiP-MS. A change in protease accessibility in this setup could either capture the direct binding site of the compound to α-Synuclein or could indicate structural changes that occur as a consequence of binding, outside of the compound binding site itself.

First, we investigated structural changes of α-Synuclein monomer upon addition of EGCG for 5 minutes. We observed changes in protease accessibility of the N-and C-termini in monomeric α- Synuclein in the presence of EGCG, primarily in regions containing aromatic residues (**Figure 4A**). Previous reports have suggested unspecific binding of EGCG to the protein backbone^40^, which we could reproduce by NMR analysis (1:10 molar ratio monomeric α-Synuclein: EGCG) (**Figure 4B**). Since our LiP-MS observations are specifically at the N-and C-termini, these could indicate an additional structural change due to the interaction with EGCG. To test this, we employed paramagnetic relaxation enhancement (PRE) NMR using A91C-α-Synuclein and (1-oxy-2,2,5,5-tetramethyl-*D*-pyrroline-3- methyl)-methanethiosulfonate (MTSL) labelling to determine changes in long-range contacts within the α-Synuclein monomer in the presence of EGCG. We observed that the ratio of MTSL / no MTSL at the N-and very C-terminus was lower in the presence of EGCG (**Figure 4C**), indicating that the interaction of EGCG with monomeric α-Synuclein induces a compaction of the protein. Truncated versions of α-Synuclein lacking the N- (Δ2-11) and the C-terminus (Δ122-140) had a greater propensity to aggregate, as previously shown^41,42^, but in comparison to the WT protein, EGCG had a smaller effect on their aggregation (**Figure 4D**). Thus, the N-and C-termini of α-Synuclein are involved in the inhibition of aggregation by EGCG further supporting our LiP-MS based observation of conformational compaction of N-and C-terminus induced by EGCG.

**Figure 4.**
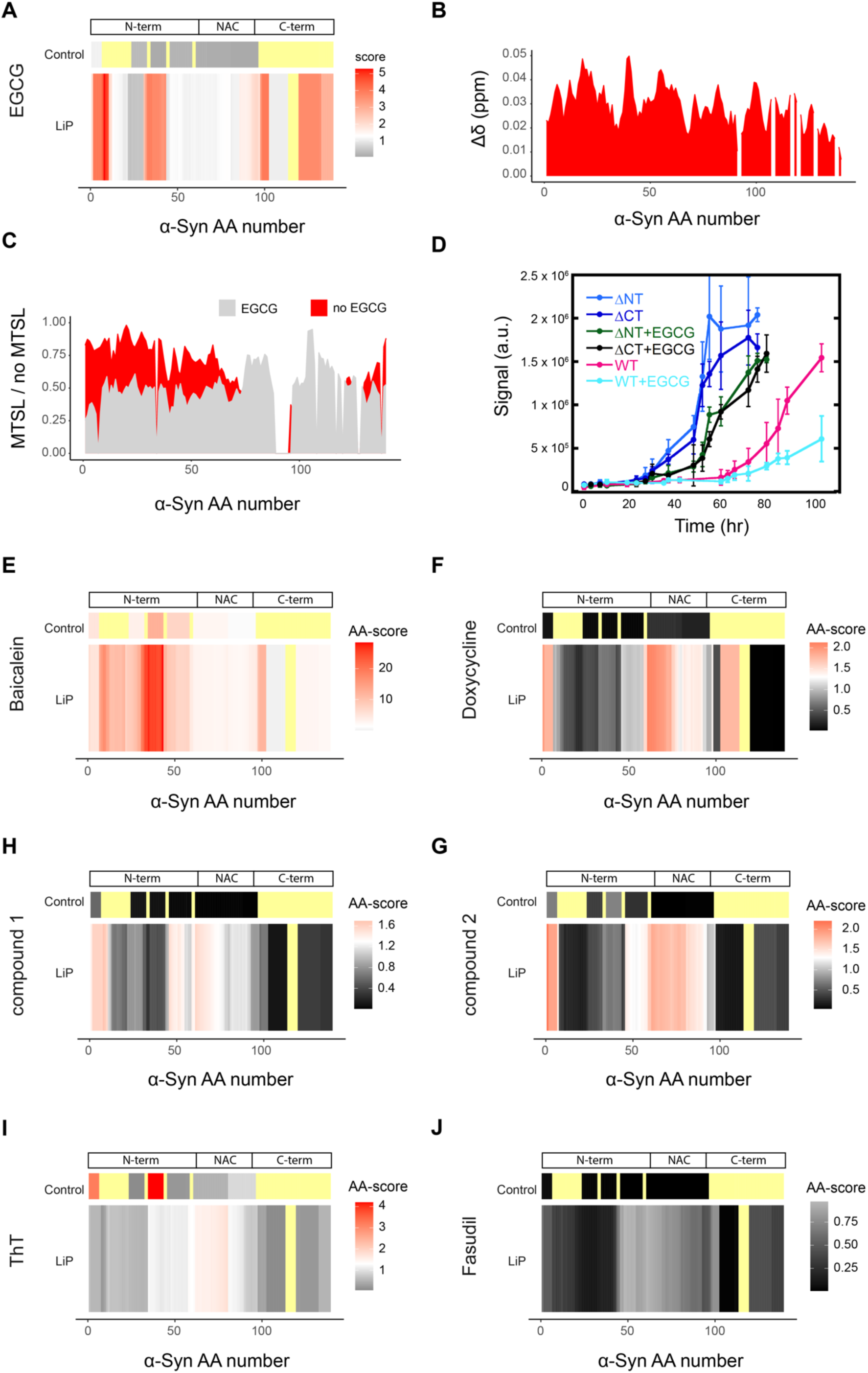
Structural changes within monomeric α-Synuclein in the presence of compounds. **A** Fingerprint of EGCG-treated α-Synuclein monomer compared to untreated α-Synuclein monomer. The upper panel indicates control (trypsin-only) peptide analysis. The lower panel indicates amino acid- level analysis of LiP peptides; the scales in both fingerprints (i.e., black to red) indicates the score per amino acid or peptide. The significance threshold of −log_10_(0.05) x log_2_(2) is shown in white, with red indicating higher scores. The more intense the red colour, the higher the score. Not significant in grey and black. Not detected in pale yellow. **B** NMR spectra of α-Synuclein with 10 x EGCG. **C** MTSL / no MTSL of no EGCG (light grey) and 10 x EGCG (red) treated α-Synuclein monomer. **D** ThT assay with ΔN (Δ2-11) − and ΔC (Δ122-140) α-Synuclein treated with 10 x EGCG. **E-J** Fingerprint of the comparison of the untreated α-Synuclein monomer structure and α-Synuclein monomers treated with Baicalein (**E**), Doxycycline (**F**), compound #2 (**G**), compound #1 (**H**), Thioflavin T (**I**) and Fasudil (**J**). Scale as in (**A**).

We assessed whether the tested compounds covalently bound α-Synuclein using data from the control condition, in which samples were digested only with trypsin, a site-specific protease, under denaturing conditions. Covalent binding is expected to yield a decrease in abundance of peptides including the covalent modification site due to the resulting mass shift. Thus, a decrease in intensity of tryptic peptides in the tryptic control condition upon addition of a compound may indicate covalent binding events. We detected no evidence of covalent binding of EGCG to α-Synuclein monomer. Baicalein however did show tryptic peptides with decreased intensity (3 peptides out of 6 detected peptides) (**Figure 4E**), likely indicating covalent modification of α-Synuclein monomer by Baicalein, as previously reported^38,43^. The parallel structural (i.e., LiP) analysis indicated that Baicalein also had structural effects on α-Synuclein monomer. Since the sites of covalent modification overlapped substantially with the observed structural changes, we normalised the peptide-level LiP data with the tryptic peptides (see Methods). The Baicalein-dependent structural changes no longer persist after this normalization (**Supplementary Figure 8**), indicating that the detected LiP changes may reflect these covalent changes alone. Unexpectedly, we also observed changes that could be consistent with covalent modification for α-Synuclein monomer treated with ThT (**Figure 4I**), however these occur at regions that do not overlap with the LiP changes.

Structural alterations of monomeric α-Synuclein in the presence of the other compounds were less pronounced than those for EGCG and Baicalein (**Figure 4F-I**). Nevertheless, we observed significant changes, in monomeric α-Synuclein but within the region that forms the fibril amyloid core, in the presence of compounds #1 and #2, and unexpectedly, in the presence of Doxycycline^44^ and ThT. Finally, we did not observe any structural change in the presence of Fasudil (**Figure 4J**), although a C-terminal interaction has been previously postulated^39^.

Overall, we showed that LiP-MS can capture structural alterations or direct interactions, including covalent binding events, of compounds with an unstructured amyloidogenic protein and thus can provide insight into their anti-amyloidogenic mechanism at the level of the monomeric protein.

### Compound interactions with fibrillar *α*-Synuclein

To assess compound interactions (i.e., direct binding or structural changes due to binding) with fibrillar α-Synuclein (**Figure 5**), we prepared α-Synuclein fibrils, confirmed the presence of amyloid fibrils by ThT emission and TEM (**Supplementary Figure 7B, C**), and compared their protease accessibility in the presence (5 minutes of incubation at RT) and absence of each compound. In the presence of EGCG, the intensities of several tryptic peptides (5 out of 6 detected peptides) decreased compared to the DMSO control condition (**Supplementary Figure 9**), indicating covalent modification of fibrils by the EGCG, as previously suggested^45^. Using LiP-MS to compare the proteolytic patterns of α-Synuclein fibrils in the presence and absence of EGCG, we observed structural changes at the N- terminus, as well as at the C-terminal end of the NAC core region, in the presence of the compound (**Figure 5A**). Interestingly, these changes overlap with previously predicted EGCG binding sites or structural changes^46,47^ (**Figure 5B**).

**Figure 5.**
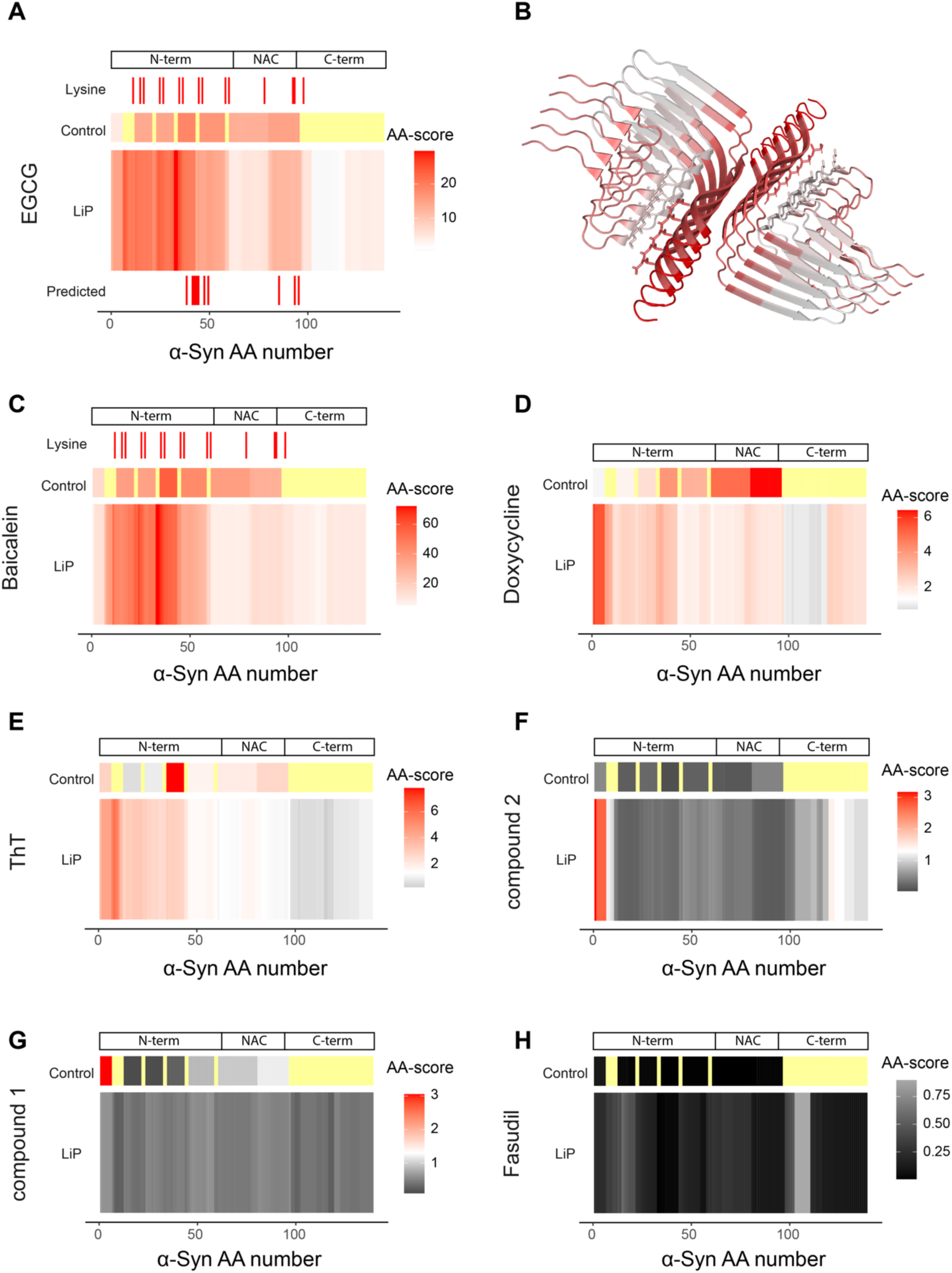
Structural changes of α-Synuclein fibrils in the presence of compounds. **A** Fingerprint of EGCG treated α-Synuclein fibril compared to untreated α-Synuclein fibril (upper panel control peptide analysis, middle panel amino acid analysis of LiP results, lower panel predicted interaction sites). The position of lysine residues is shown. The scale indicates the score per amino acid. The significance threshold of −log_10_(0.05) x log_2_(2) is shown in white, with red indicating higher scores. The more intense the red colour, the higher the score. Not significant in grey. **B** EGCG fingerprint mapped on the α-Synuclein fibril structure (pdb: 6cu7); color scheme as in A. **C-H** Fingerprint of the comparison of the untreated α-Synuclein fibril structure and α-Synuclein fibrils treated with Baicalein (**C**), Doxycycline (**D**), Thioflavin T (**E**), compound #2 (**F**), compound #1 (**G**) and Fasudil (**H**). Scale and colors as in (**A**).

In contrast to our observations with baicalein-treated α-Synuclein monomer, LiP changes in this case persisted after normalisation to the trypsin-only control peptide intensities (**Supplementary Figure 9D**), indicating both structural changes as well as covalent modifications in the affected regions. These structural changes could be a consequence of the covalent modification but may also be due to known crosslinking of fibrils by EGCG, which may affect the limited proteolysis step. Baicalein caused a similar structural response as EGCG in fibrillar α-Synuclein (**Figure 5C**) in line with molecular dynamic simulations^48^, again with changes in tryptic control peptide intensity (**Supplementary Figure 9C**) suggesting that Baicalein covalently modifies α-Synuclein fibrils as well. Notably, the other potent aggregation inhibitor, compound #2, showed a completely different interaction pattern with α- Synuclein fibrils, causing structural changes only in the extreme N terminus of the protein (**Figure 5F**).

Addition of Doxycycline caused a structural change in fibrils, consistent with previous studies that have postulated binding of Doxycycline to oligomeric species and potentially also fibrils^49^, with the most pronounced structural responses also at the very N-terminus (**Figure 5D**). Interestingly, ThT also caused structural changes at the very N-terminus of fibrils (**Figure 5E**) in regions partially overlapping with those that change in the presence of EGCG; since EGCG is known to compete with ThT binding to α-Synuclein fibrils^50^, this may reflect binding of the two molecules at the same or overlapping sites. We observed only minor changes in the amyloid core, although ThT is thought to bind to this region of the protein^51^. Finally, Fasudil and compound #1 did not yield any structural changes in fibrillar α- Synuclein suggesting that they do not interact with these structures (**Figure 5H, I**).

These data show that LiP-MS can be used to characterize structural effects of compounds on amyloid fibrils *in vitro*. LiP-MS reported EGCG interactions with α-Synuclein fibrils as predicted by molecular dynamic simulations; baicalein caused similar structural changes of the fibrils and thus likely interacts with them in a similar way. We observed an interaction of doxycycline with α-Synuclein fibrils, identified involvement of the protein N-terminus in this interaction, and report a more prominent interaction of ThT with the N-terminus than with the amyloid core of fibrillar α-Synuclein.

### In situ structural effects of anti-amyloidogenic compounds

Although *in vitro* studies allow detailed analysis of anti-amyloidogenic compound mechanisms, it is critical to assess compounds in a more physiological context. Despite extensive research, it is still not clear if *in vivo* structures of amyloid fibrils resemble those in *in vitro* models, and thus interactions with compounds in cells and tissues may be different from those *in vitro*. Further, binding of the compounds to other cellular components may result in off-target effects and reduce the anti-amyloidogenic effect due to competition or sequestration; indeed EGCG and baicalein belong to classes of molecules that are known to have promiscuous binding profiles^52–54^, and their phenotypic effects might be independent of binding to amyloidogenic protein targets. A substantial advantage of LiP-MS is that it can be applied in cell or tissue lysates and thus probe effects of compounds on the whole proteome in a near-native state.

We used LiP-MS to test the effects of Doxycycline, EGCG, Baicalein and ThT on lysates of SH-SH5Y neuroblastoma cells overexpressing α-Synuclein, which are known to form α-Synuclein inclusions^55,56^ but where the nature of these inclusions (i.e., amorphous or amyloid) is unknown. We observed structural changes in multiple proteins upon addition of each of the tested compounds (100 uM), with α-Synuclein detected as a weak hit in response to EGCG and Baicalein (log_2_FC| >1, q-value < 0.05) and showing no changes in response to the other two compounds (Figure 6A, Supp. Fig 10 A). Overall, we detected 1102, 3249, 1856, and 877 proteins showing structural changes in the cell lysate upon addition of doxycycline, EGCG, Baicalein and ThT. We went on to test the effects of EGCG on lysates of postmortem human brain, analysing cingulate gyrus pooled from two individuals with Parkinson’s disease. As in the cell lysates, we detected some α-Synuclein peptides structurally responding to EGCG, but also detected structural changes in 2489 other proteins (|log_2_FC| >1, q-value < 0.05) (**Figure 6B**). At a more stringent threshold, 949 proteins in the brain lysate showed structural changes upon addition of EGCG, but α-Synuclein was not among the hits (**Supplementary Table 1**). In both brain and cell lysate, structural changes upon addition of EGCG mapped to the NAC region of α-Synuclein (**Figure 6C**), with some additional N terminal changes apparent in the cell lysate.

**Figure 6.**
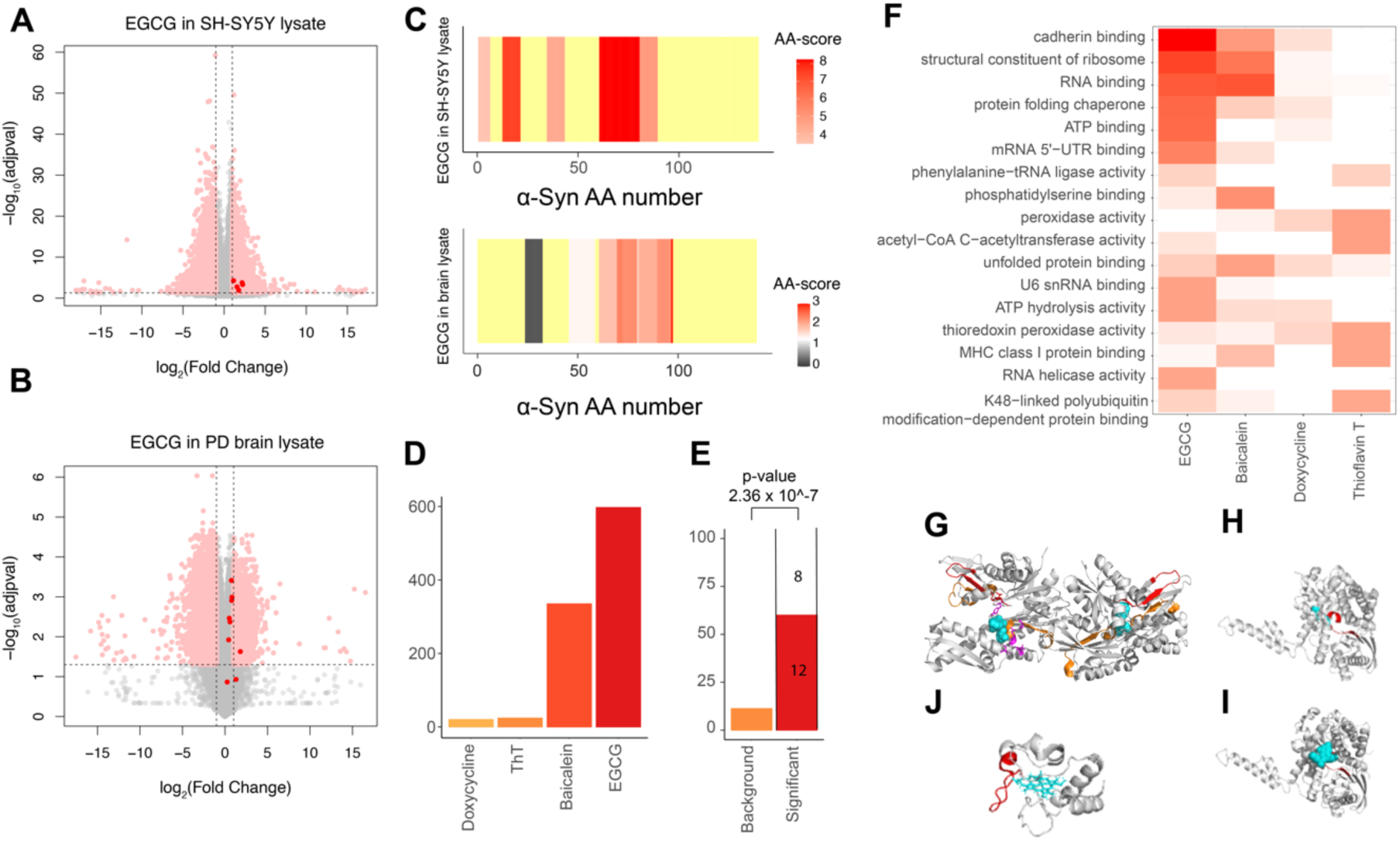
Putative in situ binding targets of anti-amyloidogenic compounds. **A, B** Volcano plots showing peptides with altered abundance after addition of EGCG to a cell lysate (**A**) and brain lysate (**B**). Dotted lines indicate the significance threshold (<u>|</u>log_2_FC| >1, q-value < 0.05; peptides from α - synuclein are coloured in red. (**C**) Structural fingerprint of α-Synuclein in cell (top) and brain (bottom) lysate. The scale indicates the score per amino acid. The significance threshold of −log_10_(0.05) x log_2_(2) is shown in white, with red indicating higher scores. The more intense the red colour, the higher the score. Not significant in grey. Not detected in yellow. (**D)**. Number of putative targets identified for the indicated compounds in a LiPQuant analysis in cell lysate (LiPQuant score > 2). (**E).** Plotted is the percentage of LiPQuant hits for doxycycline (LiPQuant score > 2) (red) or background proteome (orange) that were previously identified in a doxycycline pulldown^58^. Enrichment over background was calculated using Fisher’s exact test. (**F)** GO enrichment analysis (molecular function) for putative target proteins of the indicated compounds (LiPQuant score > 2). (**G)** Significant peptide (orange for EGCG, red for Baicalein) mapped on GRP78 structure (pdb: 3ldp). Small molecule inhibitor 3P1 in cyan. Binding site in purple. **(H)** Significant peptide (red) mapped on GLUD1 structure (pdb: 1l1f). ATP binding site extracted from “www.uniport.org” in cyan. (**I)** Significant peptide (red) mapped on bovine GLUD1 structure (pdb: 6dhl). ECG in cyan. (**J)** The two significant peptides (red) of Cytochrome C in the presence of Baicalein mapped on the Cytochrome C structure (pdb: 1j3s). HEME C in cyan. Significant peptides were defined as those with a LiPQuant score > 2.5.

Given the large number of hits in both lysates, we used our previously developed LiPQuant approach, which uses multiple criteria to define a score that can prioritize potential compound targets with high confidence^57^. We applied a dosage series of all compounds to SH-SY5Y cell lysates, restricting the analysis to cellular samples due to human brain sample availability constraints. At a stringent LiP-Quant score threshold (LiPQuant score 2.0) that we have previously used in target deconvolution experiments, we observed no evidence of α-Synuclein (55-58% coverage; C-terminus not detected) as a binding target of any of these four compounds (**Supplementary Figure 10B**). Instead, many other proteins were identified as putative targets, with multiple hits for EGCG and Baicalein (420 and 253 proteins out of 5050 and 5566 detected proteins), and fewer for ThT and Doxycycline (18 and 23 proteins out of 5650 and 4993 detected proteins) (**Figure 6D**; LiP-Quant score threshold of 2.0). At a lower LiPQuant score threshold (1.5), EGCG, but none of the other compounds, showed two significant peptides in α-Synuclein **Supplementary Figure 10C**). Binding curves for these peptides indicate a relatively low affinity interaction (**Supplementary Figure 10D**), and many other potential target proteins of EGCG (n=1550) were also identified (**Supplementary Figure 10E**). The data thus indicate that α-Synuclein is not a major binding target of EGCG, Baicalein, Doxycycline or ThT in a cell lysate and that the several other potential interactors of these compounds could compete for binding to α-Synuclein. Although we cannot rule out *in situ* binding to the protein C-terminus due to lack of coverage of this region, we consider this unlikely since we detected no strong C-terminal interaction for any compound with either the monomeric or fibrillar protein *in vitro*.

We next analysed the detected *in situ* interactors of the anti-amyloidogenic compounds in light of the literature. Previously identified direct interactors of Doxycycline^58,59^ were significantly enriched in the corresponding LiPQuant hitlist (**Figure 6E**, LiPQuant score > 2; **Supplementary Figure 10F**, LiPQuant score > 1.5), indicating that the approach captures known binding targets. EGCG is classified as pan-assay interference (PAIN) compound^59^ and is known for its promiscuity^52–54^. Consistent with this, our proteome-wide screen identified numerous potential interactors (420 proteins; **Figure 6D**, LiPQuant score > 2). Interestingly, GO enrichment analyses revealed that putative targets of EGCG were enriched for ATP binding proteins (**Figure 6F**), with 93 of the putative EGCG interactors being known ATP-binding proteins. For GRP78 (BiP), a known binder of EGCG^60^, the significantly changing LiP peptides mapped close to the predicted EGCG binding site on the protein structure (**Figure 6G**).

Similarly, changing peptides mapped close to the ATP binding site of glutamate dehydrogenase (GLUD1), known to be inhibited by EGCG^61^ (**Figure 6H)** and the changing peptide exactly maps to the known binding site of epicatechin-3-gallate (ECG) on the bovine enzyme^61^ (**Figure 6I**). Indeed, LiP-Quant-detected structural alterations of multiple ATP binding proteins (ACTR, HK1, MYH10, PAICS, ACTB) were consistently near ATP binding sites (**Supplementary Figure 11**). Interestingly, the effect of EGCG for ATP binding proteins, as for all detected interactors, was observed at median ed50 values of 25.6 μM, which indicates relatively low affinity of EGCG towards these targets. We also observed that 4 putative EGCG targets were annotated as NAD-binding proteins (e.g., ADH5, MDH2) and that these also showed significantly changing peptides mapping near NAD binding pockets (**Supplementary Figure 12**). Finally, ATP-binding proteins (169 proteins) and NAD-binding proteins (19 proteins) were also enriched (GO analysis, ‘Molecular function’; q val<0.05) among the EGCG hits within PD brain lysates, in single-dose LiP-MS experiments. Our data thus show that EGCG binds ATP- and NAD-binding pockets in complex lysates and at proteome scale. In the case of Baicalein, which is known to affect mitochondrial function^62^, complex I or complex I related components (NDUFV1, NDUFAF2) and cytochrome C were amongst the high confidence LiPQuant hits (LiPQuant score > 2.5), with the two top peptide hits mapping to a single region of Cytochrome C (**Figure 6J**). As in the case of EGCG, Baicalein showed promiscuous interactions (n=253 proteins at a LiPQuant score> 2). Putative Thioflavin T targets included several proteins (UBE2I, UBLCP1, USP14, UBQLN4, NPLOC4) in the ubiquitin-proteasome system (UPS) and related processes, which may explain its known effects on protein homeostasis^63,64^, as reflected in our GO enrichment analysis.

Overall, *in situ* LiP-MS analyses allowed the identification of known and previously unknown putative cellular interactors of anti-amyloidogenic compounds from cell lysates and pinpointed putative binding sites. The compounds under investigation bound to multiple cellular target proteins with higher affinities than to α-Synuclein or did not bind α-Synuclein in this context at all. Our data emphasize the importance of studying compound mechanisms *in situ* as well as on purified proteins.

## Discussion

We developed a modular pipeline consisting of a series of high-resolution LiP-MS analyses under different experimental conditions to study the mechanism of action of anti-amyloidogenic compounds and the amyloid binder ThT. This approach enabled us to probe whether a compound interacts with different structural forms of an amyloidogenic protein of interest, whether the compound covalently binds the protein and/or affects its aggregation, and whether it interacts with the target protein *in situ*. We applied this approach to six compounds targeting the canonical Parkinson’s disease protein α-Synuclein.

Our approach mapped changes in the amyloid core in a seeded α-Synuclein aggregation assay and these changes were sensitive to three out of six tested anti-amyloidogenic compounds. In the presence of baicalein, the green tea polyphenol EGCG and compound #2, the three most potent inhibitors of aggregation as measured by ThT fluorescence, α-Synuclein showed relatively subtle structural changes compared to the α-Synuclein monomer. Moreover, the patterns of structural change were similar for all three compounds. Since both ThT fluorescence assays and our LiP data indicated a lack of fibril formation, this could suggest a much slower rate of fibril formation or the formation of other structures, such as stable oligomers, that did not evolve to amyloid fibrils in the presence of these compounds^37,38^. Our approach also identified structure-specific effects of the compounds on purified monomeric and fibrillar forms of α-Synuclein *in vitro*. Since EGCG, baicalein and compound #2 showed different overall interaction fingerprints with α-Synuclein monomers, the data suggest that they induced similar end structures despite different interactions with monomers.

We observed an interesting discrepancy between the ThT fluorescence assay and the LiP-MS structural data for α-Synuclein aggregation in the presence of doxycycline and compound #1. For both compounds, ThT fluorescence indicated that fibril formation was substantially reduced relative to control. At the same time, the LiP data indicate that fibril structures similar to control were formed at the end of the time course. A possible explanation for these data is that doxycycline or compound #1 compete with ThT for binding to α-Synuclein, or otherwise interfere with ThT fluorescence. Indeed, ThT, compound #1 and doxycycline all interacted with the N-terminal region of the NAC in the α-Synuclein monomer. While compound #2 also showed interactions with this region, and could in principle also affect the ThT fluorescence assay, in this case the structural data were consistent with the ThT data and indicated that compound #2 prevented formation of α-Synuclein fibrils. An alternative explanation for this discrepancy is that the LiP-MS signal in the presence of doxycycline or compound #1 may simply reflect a smaller amount of α-Synuclein fibrils formed under these conditions, or a fibrillar form that does not interact with ThT.

EGCG induced an N- and C-terminus-dependent compaction of α-Synuclein monomers. Previous NMR data have suggested that the monomeric protein undergoes N-C terminal interactions in the absence of any added compound^65^, consistent also with our own previous LiP-MS data showing protection of some positions in the N-terminal region; our data now suggest that EGCG may stabilize this compacted form of the protein. This is also in line with observations from ion mobility shift MS that suggested that EGCG engages α-Synuclein in compact conformations^66^. For α-Synuclein amyloid fibrils, a structural form that poses many challenges for current methods, our LiP-MS based approach revealed that the N-terminus and the C-terminal end of the NAC are involved in the interaction with EGCG, corroborating predictions from molecular dynamic simulations. While baicalein and EGCG interact very similarly with α-Synuclein fibrils, the other potent aggregation inhibitor compound #2 has a completely different interaction fingerprint, with exclusively the first 10 residues of α-Synuclein involved in the interaction, suggesting a different molecular mechanism. Doxycycline also interacted with fibrils *in vitro*; it remains to be tested whether fibril interactions of the compounds we assayed leads to disaggregation on longer time scales.

ThT interaction with amyloids is of particular interest as the PET tracer Pittsburgh Compound B^67^, used for imaging of β-amyloid plaques, is a radioactive analogue of this compound. Further, ThT aggregation assays are the main tool used to identify inhibitors of aggregation and thus understanding its mode of interaction with amyloid fibrils is of high interest. ThT is proposed to bind the amyloid-β peptide at β-sheets either by binding to hydrophobic channels along the amyloid-β peptide fibril axis^68^ or to surfaces containing aromatic residues^69^. In both cases, the cross-β sheet structure is key for its interaction. Interestingly, ThT induced clear changes in proteolytic accessibility in the N-terminus of α-Synuclein fibrils *in vitro* but only very minor changes in the aggregation core. Several prior observations are consistent with an interaction of ThT with the α-Synuclein N-terminus. The majority of cryo-EM structures of α-Synuclein fibrils resolved to date showed that at least part of the fibril N-terminus is structured, potentially allowing ThT interactions at those sites^12,70–72^. Also there have been previous reports of ThT-negative α-Synuclein fibrils, suggesting that amyloid aggregation alone is not sufficient for ThT signal^73^. In addition, the N-terminus of α-Synuclein is positively charged, as a result of which it could electrostatically interact with negatively charged ThT. The N-terminus also contains aromatic residues, as does the amyloid-beta peptide^69^, and the peptide self-assembly mimetics (PSAMs) with which ThT interaction has been characterized^74^, while the α- Synuclein aggregation core lacks aromatic residues. Further, regions of the N-terminus have been implicated in aSyn aggregation, consistent with ThT binding to this region^75^.

Fasudil showed no evidence for interaction with α-Synuclein monomer or fibril *in vitro*, which was somewhat surprising given prior NMR data and molecular dynamics simulations^39^. However, since previous work postulated a C-terminal interaction for this compound, our results could be due to incomplete coverage of the C-terminus, in addition to differences in the experimental setups. Our data are nevertheless consistent with the fact that we saw no effect of Fasudil on α-Synuclein aggregation in our setup: the structure formed after 17h of aggregation was the same in the presence and absence of Fasudil and we also observed no change in the ThT profile due to the compound.

Although *in vitro* studies of anti-amyloidogenic compounds are valuable for detailed mechanistic understanding of how the compounds may affect the aggregation of the amyloidogenic protein of interest, *in vitro*-identified molecular events may not occur, or may be functionally irrelevant, *in vivo*. Indeed, despite easily detectable structural changes compounds were applied to purified α-Synuclein monomers or fibrils *in vitro*, none of the compounds we tested *in situ* (EGCG, Baicalein, ThT and Doxycycline) showed a strong structural effect on α-Synuclein within a complex lysate. While we could identify α-Synuclein as a relatively low affinity hit of EGCG and Baicalein in complex lysates, these compounds also had numerous other higher-affinity putative binders. We note that the cellular model we used (SH-SY5Y neuroblastoma cells overexpressing α-Synuclein) is known to form α- Synuclein inclusions^55,56^, but the nature of these inclusions (i.e., amorphous or amyloid) is unknown. Our data indicate that effects of EGCG, Baicalein, ThT and Doxycycline previously observed in cellular or animal models of neurodegeneration^76–81^ are likely due to interactions with proteins other than α- Synuclein.

There are many potential reasons for the discrepancy between the *in vitro* and *in situ* results. Given competing binders within the complex proteome, the amount of compound available for α-Synuclein interaction is likely to be much lower in the lysate than for purified protein *in vitro*. Indeed, in brain lysates, half of the top 10 EGCG hits were 1-2 orders of magnitude more abundant than alpha- synuclein, which may in part explain preferential binding to these proteins. Further, the α-Synuclein coverage *in situ* was incomplete. It is possible that the compounds interact with the α-Synuclein C-terminus, which we did not cover *in situ*, although this region did not seem important for compound interaction based on *in vitro* data. There may also be biological reasons for the discrepancy. α- Synuclein is thought to be conformationally complex, and it is possible that the structure of the protein in cell lysates is not identical to either the purified unfolded monomer or to the amyloid fibril we used in our *in vitro* experiments. Overall, our data argue strongly that studies of drug mechanism should be done also *in situ,* to uncover potential liabilities when moving into more complex systems such as animal models. Our approach will enable such studies, and importantly, will enable screening for anti-amyloidogenic drugs as well as for diagnostic agents such as PET tracers directly in brain lysates. This would target such screening efforts directly at physiologically and pathologically relevant structures, for instance by identifying compounds that selectively bind the pathological conformation of α-Synuclein in brain lysates of PD patients but have no interaction with any proteins in healthy individuals. Our approach would also allow the identification of putative off-targets in a physiologically relevant context.

We report the first proteome-wide analysis of interactors of EGCG. As a pan-assay interference (PAIN) compound, due to its propensity to interact with membranes^54^ and to covalently modify proteins^53^, EGCG is recognized as neither a good drug candidate nor suitable for SAR optimization. It is nevertheless studied as a potential drug candidate in numerous diseases such as cancer, metabolic syndrome, and neurodegeneration^52–54^, and is being tested as a drug or a dietary supplement in clinical trials for numerous conditions as well. Given relatively lax regulation for dietary supplements, EGCG and/or green tea extract is already promoted as a supplement for weight loss, heart health, inflammation and even cognitive protection, including for individuals with Down’s syndrome who are at high risk for Alzheimer’s disease. Our analysis has identified numerous potential interactors of EGCG in human cell lysates, with an enrichment for ATP-binding proteins, consistent with its known promiscuity and with prior data reporting competitive binding at ATP-binding sites of individual proteins (e.g., PI3K, mTOR, ZAP-70, glutamate dehydrogenase and GRP78)^60,61^. Especially in light of adverse effects that have been observed at higher doses^82^, our data strongly suggest that high doses of EGCG, such as those used in dietary supplements and in particular outside of medical supervision, should be avoided.

While EGCG and baicalein caused the most widespread effects of the four compounds we analyzed in situ, doxycycline and ThT also showed putative interactions with several cellular proteins. It is possible that these compounds bind functional amyloids physiologically present in cells, however, since the structural state of the identified interactors is unknown, this question would need to be resolved in future studies.

Our study has examined only a small number of known and proprietary anti-amyloidogenic and amyloid-binding compounds for a single amyloidogenic protein, α-Synuclein. The LiP-MS approach could however be extended to other compounds and proteins of interest. Identification of a compound that binds the endogenous structure of an amyloidogenic protein can be followed by *in vitro* analysis of compound mechanism of action using our amino acid-centric analysis. *In vitro* studies with multiple structural forms could also give insight into which form is present endogenously. Applied *in situ*, our versatile approach tests for binding in a physiologically relevant context, identifies off-target binders and thus assists in compound optimization and may suggest new ways to influence disease progression in neurodegeneration.

## Methods

### ThT aggregation assay

In the seeded ThT aggregation assay, 17.5 μM of monomeric α-Synuclein and 175 nM α-Synuclein seeds were incubated in aggregation buffer (50 mM Tris, 250 mM NaCl, pH 7.4) containing 3 % DMSO (Dimethyl sulfoxide (DMSO) ≥99.5% (GC); Cat# D4540-100ML (Sigma)) and 40 μM ThT (ThT (Cat# T3516-5G (Sigma)) stock solution 3 mM (in H_2_O)), in the presence of 100 μM of compounds. Compound stocks were prepared in 100 % DMSO and stored at −20°C at a concentration of 4 mM. The solution was thawed by 5 minutes sonication in a sonication bath (Elmasonic – X-tra 30 H (Elma)) prior to addition to the reaction mixture. The seeds were generated from α-Synuclein fibrils produced upon incubation at 37 °C under constant agitation at 1000 rpm in aggregation buffer (50 mM Tris, 250 mM NaCl, pH 7.4) at a final concentration of 1 mg/ml over 6 days of incubation. The fibrils were fragmented by 10 snap freezing and 1 min sonication cycles at 35 °C, aliquoted in low protein retention vials and stored at −20°C. The seed aliquots were only used once and discarded once thawed.

ThT aggregation was followed using an Infinite M200 PRO (TECAN) plate reader. The samples were prepared in a volume of 70 μl and then distributed in a 96-well plate (Plate 96F – non treated – Black Microwell S1 Cat# 237105 (Thermo-Fisher)). The plate was incubated in the Infinite M200 PRO (TECAN) plate reader at 37 °C, under constant orbital shaking with a 1.5 mm Amplitude. The ThT fluorescence was measured by fluorescence top reading with an excitation wavelength of 440 nm and an emission wavelength of 485 nm using a gain of 80 and 15 flashes per well. After 17 hours, the samples were either probed by LiP-MS, where different replicates per condition were probed, and the left-over samples after LiP-MS were pooled and snap frozen to perform the transmission electron microscopy and native PAGE experiments one day later.

### Transmission electron microscopy (TEM)

The carbon film coated copper grids were first glow discharged using 25 mA for 45 seconds with negative polarity. Samples were applied to the discharged grids applying 4 μl of 17.5 μM sample, dried with a filter plate before two consecutive washes in double deionised water followed by drying with filter paper. For staining, one drop of uranyl-acetate was added on top of the grid and used for one wash, followed by one drop and incubation for exactly 1 minute before the stain was removed with a filter paper and the grid was air dried. The TEM images of the time course were imaged using a TFS Morgagni 268 and the images of the seed preparation were imaged using a Hitachi7700.

### Assessment of unstructured monomer conformation

To assess the purity of α-Synuclein monomers, blue native PAGE was performed using NativePAGE™ Sample Prep Kit and precasted NativePAGE™ 4 to 16%, Bis-Tris gels (1.0 mm, Mini Protein Gel, 10-well). 1 to 3 ug of α-Synuclein was diluted with NativePAGE™ 4X Sample Buffer. Samples and the NativeMark™ Unstained Protein Standard were loaded into wells filled with 1 X NativePAGE™ Dark Blue Cathode buffer, containing Coomassie G-250, filled wells. Gels were run at 150 V constant in NativePAGE™ Dark Blue Cathode buffer at the Cathode and NativePAGE™ Anode buffer at the Anode for 30 minutes. NativePAGE™ Dark Blue Cathode buffer was exchanged with NativePAGE™ Light Blue Cathode buffer and the gel was run until completion at 150 V constant. Gels were fixed in fix solution (40 % methanol, 10 % acetic acid) and microwaved for 45 seconds, followed by 15 minutes shaking on an orbital shaker. The gels were then destained in destaining solution (8 % acetic acid) and microwaved for 45 seconds, followed by incubation on the orbital shaker for 15 minutes. This procedure was repeated multiple times, until the gel was destained to completion. SDS-PAGE was performed using precasted 4–12% NuPAGE™ Bis-Tris gels in NuPAGE™ MES SDS Running Buffer. 5 x Laemmli buffer was added to the samples containing 1 μg, 2 μg and 3 μg monomeric α-Synuclein. As a marker we used PageRuler Plus Prestained protein ladder. The gel was run at 80 V constant for 15 minutes, followed by 150 V constant until completion. The gels were stained using PageBlue™ Protein Staining Solution, and destaining was achieved by shaking on an orbital shaker in double deionized water.

### Monomer purification

For the ThT aggregation assay, α-Synuclein purchased from rPeptide (Cat# S-1001-4) was used. Experiments with α-Synuclein monomers and fibrils were performed using purified α-Synuclein from the laboratory. Wildtype α-Synuclein was expressed in transformed BL21 DE3 *E. Coli* upon Isopropyl- *β*-D-thiogalactopyranosid (IPTG) induction. The cells were lysed by employing an osmotic shock upon resuspending the *E. Coli cells* in a 40% sucrose Tris-buffer followed by transferring the pellet to deionized cold water. The solution was boiled for 10 min followed by centrifugation at 20’000 g at 4 °C for 20 min. α-Synuclein was then purified by Anion exchange chromatography using a HiTrap® Q FF 16/10 (from GE healthcare). The fractions containing α-Synuclein were dialyzed overnight against water and dried after aliquoting 200 ug into low binding Eppendorf tubes. The tubes were stored at −80 °C.

### Fibril generation

Fibrils were generated by incubating 1 mg/ml of α-Synuclein in PBS pH 7.4 under constant agitation at 800 rpm at 37 °C in a thermocycler.

### ThT aggregation of ΔN and ΔC α-Synuclein

Around 10 mg Solid lyophilized α-Synuclein was taken and dissolved in 1 ml PBS, 7.4, 0.01% sodium azide by adding few ul of 1(M) NaOH. The pH was brought back to 7.4 by adding few ul 1(N) HCl. The protein solution was centrifuged at 14’000 g for 30 minutes at 4 °C and loaded on size exclusion column to isolate the monomers. Concentration was determined by absorbance at 280 nm, considering the molar absorptivity (ε) is 5960 for α-Synuclein and all N-terminal truncated, K mutants and 1490 for C-terminal truncated mutants. Final concentration of the WT α-Synuclein and α-Synuclein mutants were adjusted to 300 μM for the aggregation studies.

1 mM ThT was prepared in Tris-HCl buffer, pH 8.0, 0.01% sodium azide. 2 μl of 1 mM ThT solution was added to the 7.5 μM protein solution (300 uM stock solution) in 200 μl PBS buffer, pH 7.4, 0.01% sodium azide. ThT fluorescence assay measurements were done using Horiba-Jobin Yvon (Fluomax4). The excitation was set to 450 nm and the emission was measured in the range of 460-500 nm. The slit width for both excitation and emission were kept at 5 nm. WT α-Synuclein and different mutant, C-terminal truncation (1-121 amino acid residues) and N-terminal truncation (2-11 amino acid residues absent from 140 amino acid sequences) were used for the aggregation studies. The concentrations of α-Synuclein was determined by absorbance at 280 nm, considering the molar absorptivity (ε) is 5960 for α-Synuclein and all N-terminal truncated, K mutants and 1490 for C-terminal truncated mutants.

### Expression and purification of ^15^N labeled N-terminal acetylated A91C-alpha Synuclein

The following plasmids were co-expressed in *E. coli* BL21*DE3 cells. Human α-Synuclein with a A91C point mutation cloned into pRK172 containing ampicillin resistance and the two N-alpha-acetyltransferase (NatB) complex subunits *naa20^+^*(SPCC16C4.12) and *naa25^+^*(SPBC1215.02c) from *S. pombe* cloned into pACYCduet containing chloramphenicol resistance^315^.

The transformed cells were grown in LB medium containing the appropriate antibiotics at 37°C up to an *A*_600_ (absorbance at 600 nm) of 1.0. The cells were transferred to M9 minimal medium containing 1g/L Ammonium-^15^N-chloride. Protein expression was launched using 1mM isopropyl-β-D-thiogalactopyranoside and continued shaking at 37°C over night.

The cells were lysed using osmotic shock by resuspending the cells in a 40% sucrose Tris-buffer and transferring the cell pellet to deionized cold water. The supernatant was boiled for 10 minutes and Ammonium sulfate was added to get a 35% saturated solution. All precipitated parts were discarded and α-Synuclein was precipitated by increasing the Ammonium sulfate content to 55% saturated. The α-Synuclein pellet was dissolved in Tris-Buffer and dialyzed. α-Synuclein was further purified by Anion exchange chromatography using a HiTrap® Q FF 16/10 (from GE healthcare). Exclusively monomeric α- Synuclein was obtain by size exclusion chromatography using a HiLoad® 26/600 Superdex® 75 prep grade (from GE healthcare). The purity was checked using SDS PAGE electrophoresis. The purified α- Synuclein was lyophilized and stored at −20°C.

### Labeling of A91C-α-Synuclein with MTSL

Lyophilized ^15^N labeled N-terminal acetylated A91C-α-Synuclein was dissolved in phosphate buffer saline at pH 7.4. Dithiothreitol (DTT) was added in a ten fold excess in order to reduce existing disulfide bridges of Cysteine 91. The protein was incubated at 10°C for 30min, followed by DTT removal using a PD-10 desalting column with Sephadex G-25 resin (from GE healthcare). The reduced protein was mixed with a ten fold excess of the nitroxide spin label MTSL and incubated at room temperature protected from light for one hour. The excess MTSL was removed using a PD-10 desalting column. To remove higher molecular α-Synuclein species after MTSL-labeling, the proteins were filtered using a 100-kDa molecular weight cut-off concentrator (Amicon). The final concentration of MTSL labeled A91C- α-Synuclein was measured using a JASCO V-650 UV-VIS spectrophotometer.

### NMR Spectroscopy

A Bruker Avance III HD 600 MHz spectrometer equipped with a triple resonance cryo probe was used for all the NMR experiments. All the NMR samples contained 50μM uniformly ^15^N labeled A91C- α- Synuclein (with or without MTSL label) in PBS pH 7.4 with 10% D2O (v/v). To perform paramagnetic relaxation enhancement analysis, all the 1H-15N HMQC spectra were acquired at 283K using Bruker Topspin 3.2 for 1 hour, with 1024 x 128 complex points with 24 scans and an interscan delay of 0.5 seconds. NMR titration experiments with Epigallocatechin gallate (EGCG) was performed by varying the concentration to be 0μM, 50μM, 150μM, 300μM and 500μM. Each sample of this titration was prepared fresh from stock solutions 30min before the measurement. All samples had a total volume of 400μl and were transferred into a 5mm Shigemi tube.

NMR spectra were processed using Bruker Topspin 4.0.6 and analyzed in NMRFAM-Sparky 1.412^316^ for visualization and peak intensity analysis. The NMR signal intensity ratios (*I*_MTSL_/*I*_0_) were determined residue-wise by dividing the maximal peak height of the 1H-15N HMQC cross peak for MTSL labeled A91C-α-Synuclein (*I*_MTSL_) through the maximal peak height of the corresponding cross peak for A91C- α-Synuclein without spin label (*I*_0_). Only spectra with equal concentration of EGCG for MTSL labeled or not spin labeled A91C- α-Syn were used to calculate a *I*_MTSL_/*I*_0_ intensity ratio. The intensity ratios (*I*_MTSL_/*I*_0_) were determined for each EGCG concentration individually, plotted as rolling average and compared with each other. Prolines, since they do not have an amid proton, and other amino acids for which the *I*_0_ values did not exceed noise level were excluded from the rolling average and no value was plotted.

### SH-SY5Y α-Synuclein overexpressing cell line generation and cell culture

Polyclonal stable α-Synuclein overexpression SH-SY5Y cell lines were produced by using lentiviral vectors as described in Francesca Macchi et al^56^. 2015, and provided by AC Immune. Overexpression was confirmed by Western Blot (*data not shown here*). Cells were cultured in DMEM-F-12/10%FBS/1% Penicillin-Streptomycin and α-Synuclein overexpression was maintained by selection with puromycin in a final medium concentration of 1 ug/ml every three weeks. Specifically, always one passage before harvesting of the pellets, the cells were selected using 1 ug/ml of puromycin dihydrochloride (ThermoFisher), then grew until 80-90 % confluency, expanded and harvested. Harvesting was done by scraping the cells of the dish, removing the medium and two washes with PBS pH 7.4. The pellets were then centrifuged in 1.5 ml Eppendorf tubes at 1000 x g at 4 °C for 5 minutes, snap frozen in liquid nitrogen and stored at −80 °C until use.

### Native protein extraction

Native protein extraction was done by resuspending pellets of α-Synuclein overexpressing SH-SY5Y cells in 200 μl of LiP-buffer (100 mM HEPES, 150 mM KCl, 1 mM MgCl_2_, pH 7.4), followed by 10 consecutive douncing steps using a pellet pestle on ice. The samples were centrifuged at 1’000 x g for 5 minutes at 4 °C to get rid of the cell debris. Patient’s brain homogenates were generated by dilution at 20% (weight:volume) and sonication in 150mM KCl, 50mM Tris-HCl pH7.5 buffer. α-Synuclein overexpressing SH-SY5Y cell extracts were directly processes by LiP-MS, patient brain homogenates were stored at −80 °C prior to LiP-MS.

### LiP-MS

For the samples prepared in the aggregation assay, 20 μl of 0.25 mg/ml (17.5 μM) α-Synuclein was used to reach a final amount of 5 μg of α-Synuclein per replicate. The volume was adjusted to 50 μl using LiP-buffer (100 mM HEPES, 150 mM KCl, 1 mM MgCl_2_, pH 7.4). In the case of purified α-Synuclein monomer, samples were freshly resuspended from lyophilised powder and additionally ultracentrifuged at 50’000 x g. The supernatant was used for the monomer fraction. In the case of the fibril, fibrils were directly taken from the thermoshaker and ultracentrifuged at 50’000 x g at 4 °C, and the pellet fraction was resuspended in LiP buffer. Protein concentrations of α-Synuclein monomers and fibrils were determined using bicinchoninic acid assay (BCA). Then 1.4 μM of α-Synuclein (1 μg in 50 μl) was incubated with 140 μM compound (1:100 molar ratio) or dimethylsulfoxid (DMSO) for exactly 5 minutes at 25 °C in a thermocycler (Biometra TRIO) before proteinase K (Proteinase K, Tritirachium album, 10 mg, Sigma Aldrich) digestion. The final DMSO concentration was 2 %.

Protein concentrations of α-Synuclein overexpressing SH-SY5Y and patient brain samples was determined using bicinchoninic acid assay (BCA). α-Synuclein overexpressing SH-SY5Y extracts were diluted to 1 mg/ml protein concentration, using a volume of 50 μl, thereby 50 ug of protein per replicate. Protein concentrations of two different patient brain samples were determined using bicinchoninic acid assay (BCA) and the same amount of sample per patient was pooled to a master mix followed by protein concentration adjustment to 1 mg/ml, using a volume of 50 μl, thereby 50 μg of protein per replicate.

α-Synuclein overexpressing SH-SY5Y replicates were incubated with dimethylsulfoxide (DMSO), 0.01 μM, 0.1 μM, 1 μM, 10 μM, 25 μM, 50 μM and 100 μM compound for exactly 5 minutes at 25 °C in a thermocycler (Biometra TRIO) before proteinase K (Proteinase K, Tritirachium album, 10 mg, Sigma Aldrich) digestion. Patient brain replicates were incubated with dimethylsulfoxide (DMSO) or 100 μM EGCG for exactly 5 minutes at 25 °C in a thermocycler before proteinase K digestion. The final DMSO concentration was 2 %.

Proteinase K was added in a 1/100 enzyme to substrate ratio in a volume of 5 μl and homogenised by pipetting up and down for exactly 20 times using a 10 μl multichannel pipette. Samples were digested for 5 minutes at 25 °C in a thermocycler (Biometra TRIO) followed by boiling for 5 minutes at 99 °C and cooled down to 4 °C. After cooling down to 4 °C for exactly 2 minutes the samples were diluted in 55 μl of a freshly prepared 10 % sodium deoxycholate solution reaching a final DOC concentration of 5 %. Samples of the ThT aggregation assay were frozen at this stage for 24 hours before further processing, whereas α-Synuclein monomer and α-Synuclein fibrils, patient brain samples and α-Synuclein overexpressing SH-SY5Y samples, were directly processed further.

### Trypsin / lysC digestion

Samples were reduced with tris(2-carboxyethyl)phosphine hydrochloride (TCEP) by adding TCEP in a final concentration of 5 mM and incubating the samples at 37 °C for 45 minutes. After reduction, samples were alkylated using a final concentration of 40 mM iodoacetamide (IAA) and incubation for 15 minutes at room temperature in the dark. The samples were then diluted to 1 % DOC using 10 mM ammonium bicarbonate (Ambic). Lysyl endopeptidase LysC (Wako Chemicals) and sequencing-grade porcine trypsin (Promega) were added in a 1/100 enzyme to substrate ratio to digest the samples in a 96-well plate at 37 °C under constant agitation at 200 rpm overnight. After overnight digestion, digestion was stopped, and DOC was precipitated by adding 50 % (vol/vol) formic acid (FA) (Carl Roth GmbH) to a final concentration of 2 %. Finally, DOC was removed by filtration using a 0.2 um PVDF membrane filter (Corning FiltrEX 96-well White Filter Plate). Filtration was done by centrifugation at 800 x g.

### Sample desalting procedure

For sample desalting, a Harvard Apparatus 96-well C18 Micro-Spin column plate was used. The wells were washed with 200 μl methanol (Carl Roth GmbH) followed by two washing steps with 200 μl buffer A (0.1 % FA). Sample were loaded and washed twice with 200 μl buffer A. Finally, samples were eluted in 50 μl of buffer B (80 % acetonitrile (ACN) in 0.1 % FA) and heat dried. To prepare the samples for mass spectrometric acquisition, the samples were resuspended in buffer A, containing iRT peptides (iRT kit, Biognosys). For data-dependent acquisitions (DDA) library generation, each replicate of each condition was pooled into one sample In the case of purified proteins, only DIA samples were measured and analyzed using directDIA 2.0.

### Instrumentation and MS data acquisition

#### Liquid chromatography

α-Synuclein overexpressing SH-SY5Y samples were measured on Orbitrap Fusion™ Lumos™ Tribrid™ mass spectrometer (ThermoFisher Scientific) and brain samples on Orbitrap Exploris mass spectrometer (ThermoFisher Scientific). For nanoelectro spray ionization (nESI), the instrument was connected to a Nanoflex electrospray source. To separate the peptides, a nano-flow LC system (Easy-nLC 1200, Thermo Fisher Scientific) and PepMap RSLC column (250 mm × 75 μm, 2 μm particle size, ThermoFisher Scientific) were used. Specifically, peptide separation was achieved by a linear gradient of lc-buffer A (5 % ACN, 0.1 % FA, Carl Roth GmbH) and lc-buffer B (95 % ACN, 0.1 % FA, Carl Roth GmbH) increasing from 3 % to 35 % lc buffer-B for 120 minutes with a flow rate of 300 nl/minute.

The other samples were measured on Orbitrap Fusion™ Lumos™ Tribrid™ mass spectrometer (Thermo Fisher Scientific). For nanoelectro spray ionization (nESI), the instrument was connected to a nano electrospray ion source. To separate the peptides, an ultra-performance liquid chromatography (UPLC) system (ACQUITY UPLC M-Class, Waters) and self-packed 40 cm x 0.75 mm i.d. columns (New Objective) containing 1.9 μm C18 beads (Dr. Maisch Reprosil-Pur 120) were used. Specifically, peptide separation was achieved by a linear gradient of lc-buffer A (5 % ACN, 0.1 % FA, Carl Roth GmbH) and lc-buffer B (95 % ACN, 0.1 % FA, Carl Roth GmbH) increasing from 3 % to 35 % lc-buffer B over 120 min for patient brain samples, respectively 60 min for purified protein samples with a flow rate of 300 nl/minute. Finally, the column was washed for 5 minutes in 90 % lc-buffer B, to avoid contamination in the next measured samples.

#### Data-dependent acquisition

Only samples of α-Synuclein overexpressing SH-SY5Y and patient brain samples were measured in data-dependent acquisition (DDA). The two experiments were measured with different instruments.

Data-dependent acquisition (DDA) of the α-Synuclein overexpressing SH-SY5Y samples was performed using the following settings. MS1 spectra were acquired over a mass range of 350-1150 m/z. The orbitrap resolution was set to 120’000. A normalized automated gain control (AGC) target of 200 % or a maximal injection time of 54 ms was used. Precursor ions with intensities above 50’000 and charge states between 2 and 7 were selected for MS/MS scans. The selected precursor ions were isolated with a quadrupole. The isolation window of the quadrupole was 1.6 m/z. After single occurrence, precursor ions were dynamically excluded (dynamic exclusion) for 60 seconds. The mass tolerance was set to 10 ppm. To fragment the precursors, high-energy collision induced dissociation (HCD) was used. The collision energy was fixed at 30 %. The MS/MS spectra were recorded on an orbitrap. The orbitrap resolution in MS/MS scans was set to 30’000. The fragment ions were measured in a scan range of 150-2000 m/z. A normalized automated gain control (AGC) target of 200 % or a maximal injection time of 54 ms was used.

Data-dependent acquisition (DDA) of the brain samples was performed using the following settings. MS1 spectra were acquired over a mass range of 350-1400 m/z. The orbitrap resolution was set to 120’000. An automated gain control (AGC) target of 8.0e5 or a maximal injection time of 54 ms was used. Precursor ions with intensities above 50’000 and charge states between 2 and 7 were selected for MS/MS scans. The selected precursor ions were isolated with a quadrupole. The isolation window of the quadrupole was 1 m/z. After single occurrence, precursor ions were dynamically excluded (dynamic exclusion) for 20 seconds. The mass tolerance was set to 10 ppm. To fragment the precursors, high-energy collision induced dissociation (HCD) was used. The collision energy was fixed at 30 %. The MS/MS spectra were recorded on an orbitrap. The orbitrap resolution in MS/MS scans was set to 30’000. The fragment ions were measured in a scan range of 150-2000 m/z. An automated gain control (AGC) target of 1.0e5 or a maximal injection time of 54 ms was used.

#### Data-independent acquisition

Data-independent acquisition (DIA) of brain samples was performed using the following settings. A mass range of 350-1400 m/z was used for MS1 survey scans. The orbitrap resolution was set to 120’000. A normalized automated gain control (AGC) target of 50 % or a maximal injection time of 100 ms was used. DIA scans were performed in 41 variable-width isolation windows. The isolation of precursor ions was done using a quadrupole. Precursor ions were fragmented by high-energy collision induced dissociation (HCD). DIA-MS/MS spectra were recorded using an orbitrap with a resolution of 30’000 and a scan range of 150-2000 m/z. The maximal injection time was set to 54 ms.

Data-independent acquisition (DIA) of purified was performed using the following settings. A mass range of 350-1400 m/z was used for MS1 survey scans. The orbitrap resolution was set to 120’000. A normalized automated gain control (AGC) target of 50 % or a maximal injection time of 100 ms was used. DIA scans were performed in 20 variable-width isolation windows. The isolation of precursor ions was done using a quadrupole. Precursor ions were fragmented by high-energy collision induced dissociation (HCD). DIA-MS/MS spectra were recorded using an orbitrap with a resolution of 30’000 and a scan range of 150-1800 m/z. The maximal injection time was set to 54 ms.

Data-independent acquisition (DIA) of the α-Synuclein overexpressing SH-SY5Y samples was performed using the following settings. A mass range of 350-1400 m/z was used for MS1 survey scans. The orbitrap resolution was set to 120’000. An automated gain control (AGC) target of 8.0e5 or a maximal injection time of 100 ms was used. DIA scans were performed in 41 variable-width isolation windows. The isolation of precursor ions was done using a quadrupole. Precursor ions were fragmented by high-energy collision induced dissociation (HCD). DIA-MS/MS spectra were recorded using an orbitrap with a resolution of 30’000 and a scan range of 150-2000 m/z. The maximal injection time was set to 54 ms.

#### Search engines

Hybrid libraries from DIA and DDA data were created by Pulsar search in Spectronaut 14. Compared to the default settings, specificity was set to semi-specific for Trypsin/P, and the minimal peptide length was set to 6 amino acids. Apart from those adjustments, default settings were used. For targeted data extraction of DIA files of patient brain samples, the default settings of Spectronaut 14 were used with a peptide level FDR of 1%. Targeted data extraction of α-Synuclein overexpressing SH-SY5Y samples was done according to recommendations of Piazza and Beaton et al. 2020^57^. In the case of purified proteins DIA data was searched directDIA 2.0 in Spectronaut 14.

#### Data analysis

In purified protein and patient brain sample data the fragment group quantity was selected for peptide precursor abundance comparison using a moderated t-test and Benjamini-Hochberg adjustment after median normalisation. The analysis was performed using R version 4.1.2 and the protti package^83^. Calculations of the amino acid scores were done by assigning the score of −log_10_(q-vale) x absolute(log_2_(fold change)) to every peptide. Amino acids were grouped and the mean score per amino acid was calculated. Where relevant (i.e., for removal of covalent modification effects), normalisation to tryptic control was done by subtraction of control intensities followed by the addition of the median intensities per peptide, prior to abundance comparison.

Data from α-Synuclein overexpressing SH-SY5Y samples was fed into the LiPQuant algorithm as described in Piazza and Beaton et al^57^. 2020. GO enrichment analysis was done using the protti function “calculate_go_enrichment”.

#### Visualization

For the fingerprint visualisation the ggplot2 package in R version 4.1.2 was used. To project the amino acid fingerprints and the peptide level fingerprints, the protti functions “find_peptide_in_structure” and “map_peptides_on_structure” were used. All the plots were exported to .pdf files from R and imported into Adobe Illustrator (Version 23.1) for the presented figures.

## Acknowledgements

We acknowledge Steve Gentleman from Imperial College London, the patients and their caregivers for their participation and contributions to this study. We thank Michele Vendruscolo (University of Cambridge) for insightful discussions. P.P. was funded by a Personalized Health and Related Technologies (PHRT) grant (PHRT-506), a Sinergia grant from the Swiss National Science Foundation (SNSF grant CRSII5_177195), the Peter Bockhoff Stiftung and the ETH Zurich foundation, Parkinson Schweiz, the EMPIRIS Foundation grant (2022-FS-353), the Synapsis Foundation - Alzheimer Research Switzerland ARS, The European Research Council (866004), and the EPIC-XS Consortium (823839), the last two under the EU Horizon 2020 program.

## Supplementary Figures

**Supplementary figure 1.**
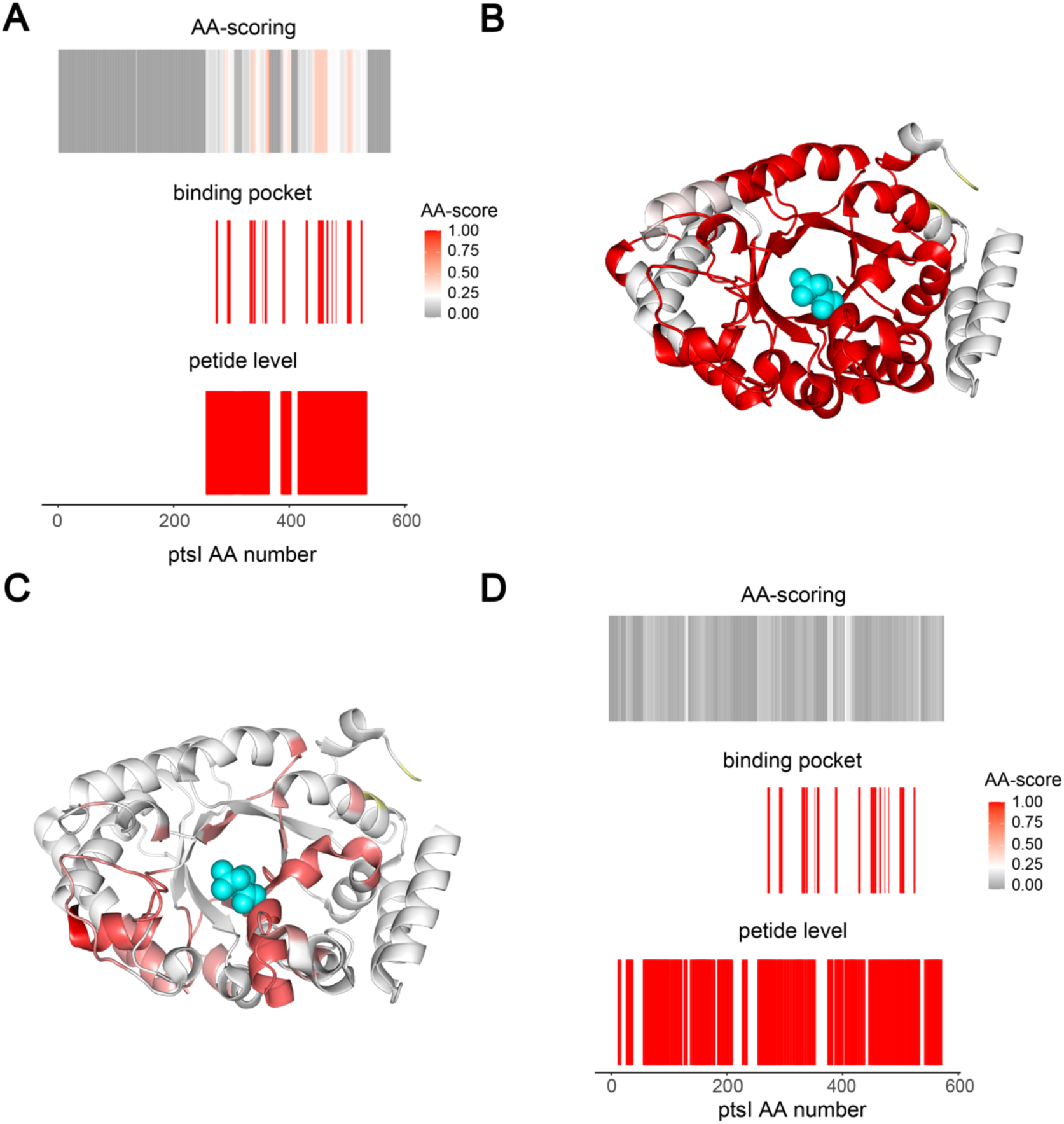
*In silico* LiP-MS of FBP-ptsI binding. To assess whether the achieved resolution of the amino acid centric analysis in FBP-ptsI binding was dependent on the specific peptides identified in this experiment, we performed an *in silico* LiP-MS experiment. We computed all potentially detectable peptides of ptsI upon proteinase K and trypsin digestion, with a minimal length of six amino acids and maximally one missed trypsin cleavage. Next, we defined the FBP binding pocket to include all atoms in a sphere of 10 Å radius around PEP. If a computed LiP peptide mapped to the binding pocket, we assigned it a score of 1, otherwise a score of 0. Peptides with a score of 1 are expected to change in a LiP-MS experiment. In an in silico amino acid-level analysis, we then averaged the assigned values per amino acid position. To test the sensitivity of the peptide-level and amino acid-level approaches to false positives, we assigned 5 % of the computed LiP peptides, picked randomly, as significantly changing (score of 1). **A** In silico amino acid centric fingerprint (top), binding pocket (middle) and peptide centric fingerprint (bottom) aligned along the ptsI sequence. **B** *in silico* peptide centric fingerprint mapped on the ptsI structure (PDB: 2xz7). **C** *in silico* amino acid centric fingerprint mapped on the ptsI structure (PDB: 2xz7). **D** Control *in silico* amino acid centric fingerprint (top)and peptide centric fingerprint (bottom) aligned along the ptsI sequence, using random peptides (5% of total peptides) assigned as significantly changing. Binding pocket defined as in A.

**Supplementary figure 2.**
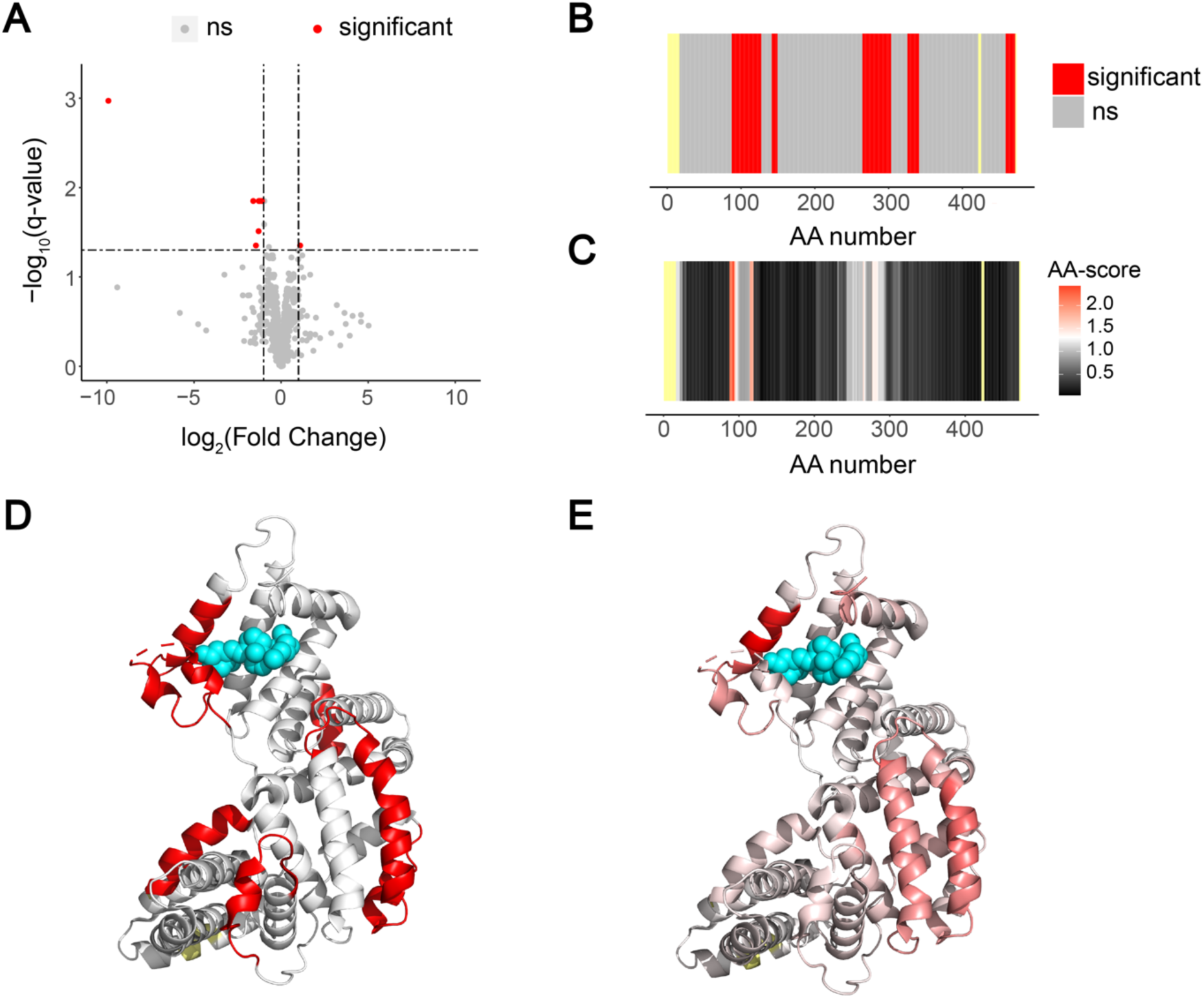
Vitamin D binding towards Vitamin D binding protein (GC). **A** volcano plot comparing peptide abundances of peptides generated in Vitamin D bound and unbound GC. **B** fingerprint of the classical LiP-MS data analysis pipeline. Significant regions in red, not significant in grey, not detected in yellow. **C** fingerprint upon scoring changes per amino acid. The scale indicates the score per amino acid. The significance threshold of −log_10_(0.05) x log_2_(2) is shown in white, with red indicating higher scores. The more intense the red colour, the higher the score. Not significant in grey. Not detected in yellow. **D** significant peptides mapped on the vitamin D binding protein structure (PDB: 1j78). Calcifediol in cyan. **E** significant amino acids of amino acid centric analysis mapped on the vitamin D binding protein structure (PDB: 1j78). Calcifediol in cyan.

**Supplementary figure 3.**
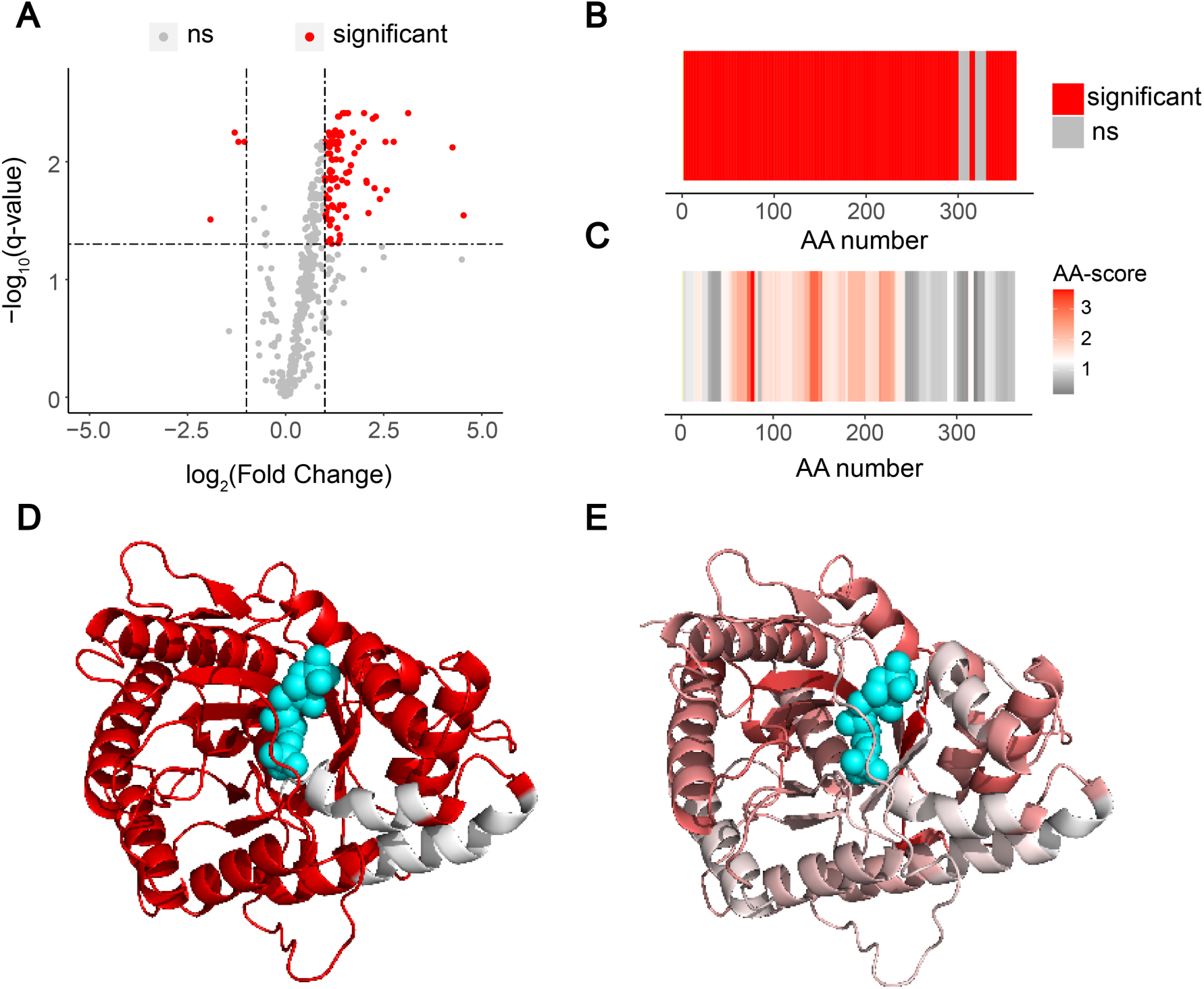
Fructose bisphosphate binding towards Fructose-bisphosphate aldolase A (ALDOA). **A** volcano plot comparing peptide abundances of peptides generated in fructose bisphosphate bound and unbound ALDOA. **B** fingerprint of the classical LiP-MS data analysis pipeline. Significant regions in red, not significant in grey, not detected in yellow. **C** fingerprint upon scoring changes per amino acid. The scale indicates the score per amino acid. The significance threshold of −log_10_(0.05) x log_2_(2) is shown in white, with red indicating higher scores. The more intense the red colour, the higher the score. Not significant in grey. Not detected in yellow. **D** significant peptides mapped on the ALDOA protein structure (PDB: 4ald). Fructose bisphosphate in cyan. **E** significant amino acids of amino acid centric analysis mapped on the ALDOA protein structure (PDB: 4ald). Fructose bisphosphate in cyan.

**Supplementary figure 4.**
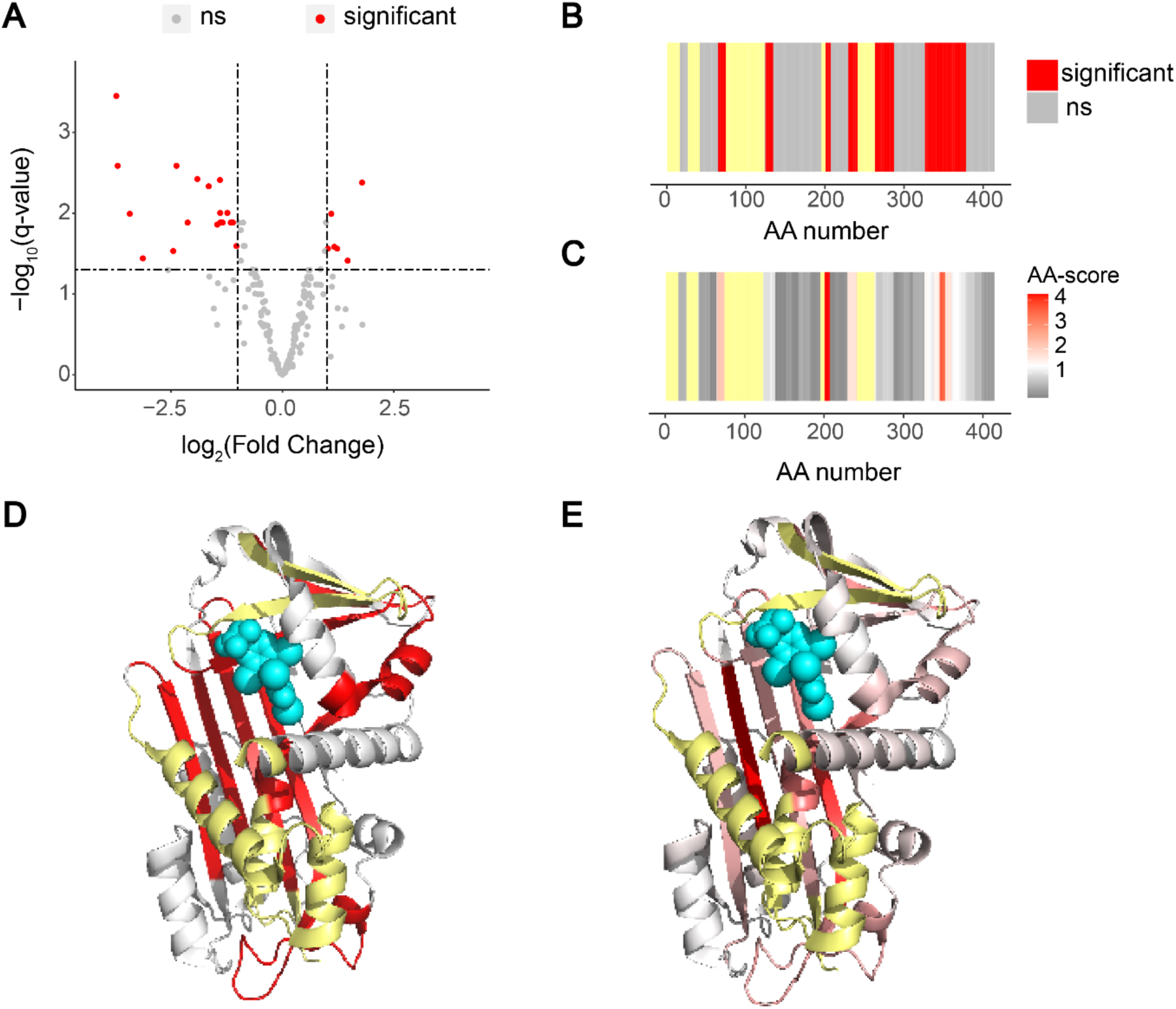
Thyroxine binding towards Thyroxine Binding Globulin (TBG). **A** volcano plot comparing peptide abundances of peptides generated in thyroxine bound and unbound TBG. **B** fingerprint of the classical LiP-MS data analysis pipeline. Significant regions in red, not significant in grey, not detected in yellow. **C** fingerprint upon scoring changes per amino acid. The scale indicates the score per amino acid. The significance threshold of −log_10_(0.05) x log_2_(2) is shown in white, with red indicating higher scores. The more intense the red colour, the higher the score. Not significant in grey. Not detected in yellow. **D** significant peptides mapped on the TBG protein structure (PDB: 2riw). Fructose bisphosphate in cyan. **E** significant amino acids of amino acid centric analysis mapped on the TPG protein structure (PDB: 2riw). Fructose bisphosphate in cyan.

**Supplementary figure 5.**
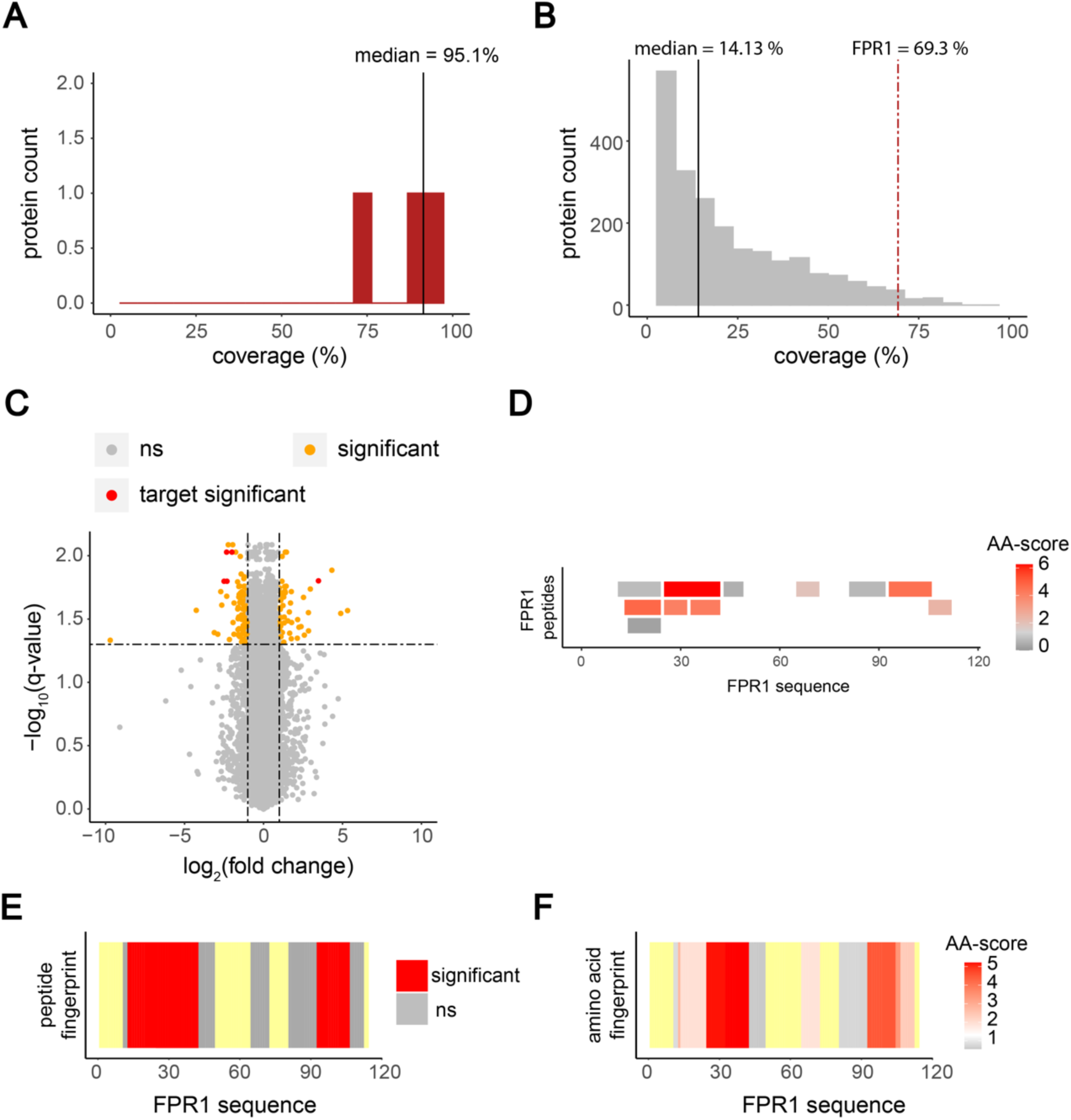
Amino acid centric analysis on a proteome wide scale. **A** Coverage of purified proteins analysed with *in vitro* LiP-MS (bin size of 5%). Median coverage is indicated by the black line (95.1%). **B** Coverage distribution of the 2417 identified proteins (bin size of 5%). Median coverage indicated by the black line (14.13 %). Coverage of FPR1 (69.3%) indicated by the red dotted line. **C** Volcano plot of rapamycin treated and untreated *S. cerevisiae* cell extracts (non-significant in light grey, significant in orange, significant and FPR1 in red). **D** Peptides and their corresponding scores (-log_10_(q- value) x absolute (log_2_(fold change)) mapped along the FPR1 sequence. Score values from grey to red. **E** Peptide-centric analysis showing significant peptides mapped along the sequence of FPR1 and onto the FK-506 bound FPR1 structure (PDB: 1yat). FK-506 in cyan. **F** Fingerprint upon amino acid centric analysis mapped along the sequence of FK-506 bound FPR1 (The scale indicates the score per amino acid. The significance threshold of −log_10_(0.05) x log_2_(2) is shown in white, with red indicating higher scores. The more intense the red colour, the higher the score. Not significant in grey.) and onto the FK- 506 bound FPR1 structure (PDB: 1yat). FK-506 in cyan.

**Supplementary figure 6.**
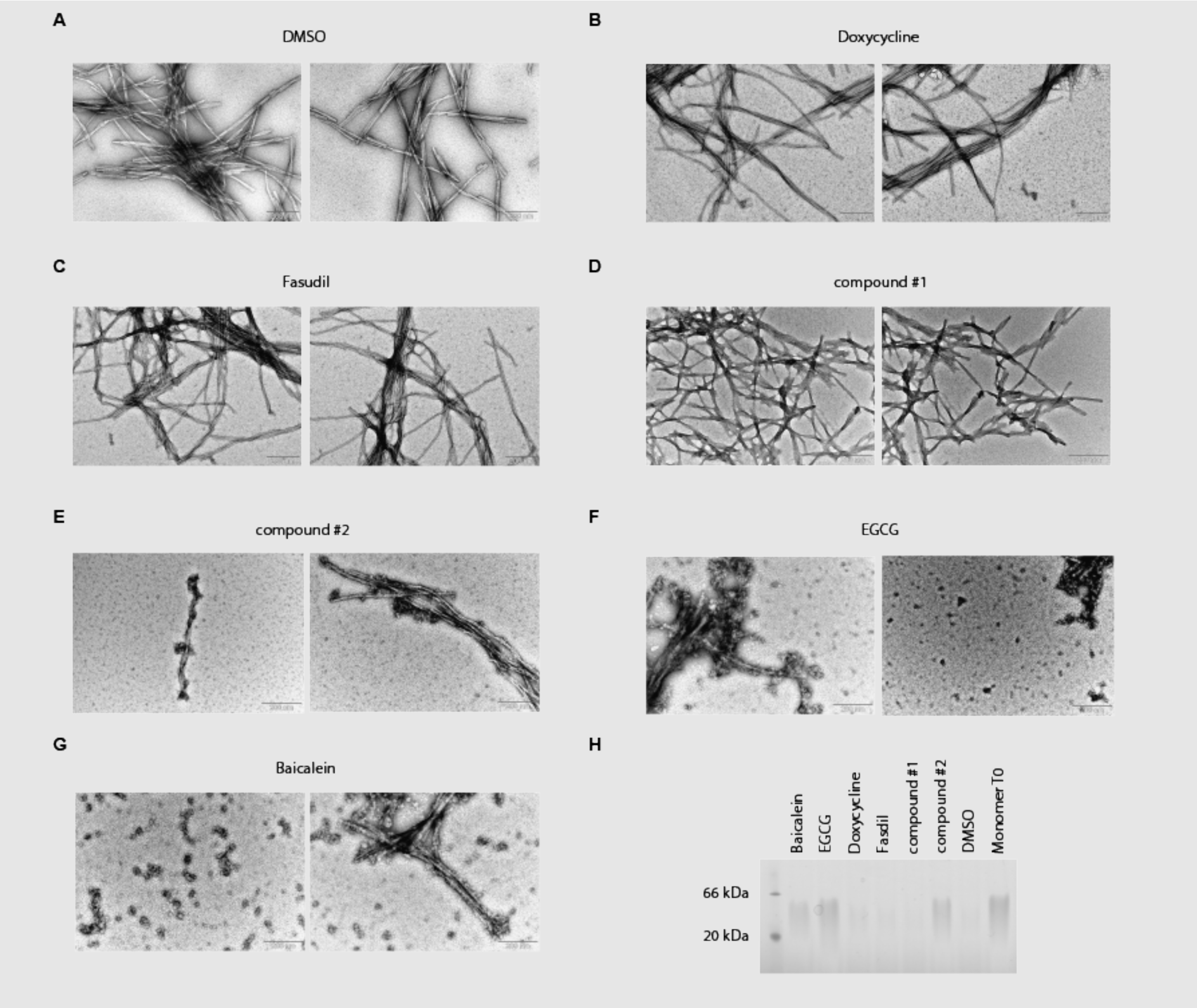
Transmission electron microscopy images of structures formed in the presence of compounds or DMSO after 17 hours of incubation. **A** DMSO control condition. **B** Doxycycline. **C** Fasudil. **D** compound #1. **E** compound #2. **F** EGCG. **G** Baicalein. **H** Native PAGE after 17 hours of incubation in the indicated conditions, compared to monomer at time point 0. Oligomeric and fibrillar forms are not resolved.

**Supplementary figure 7.**
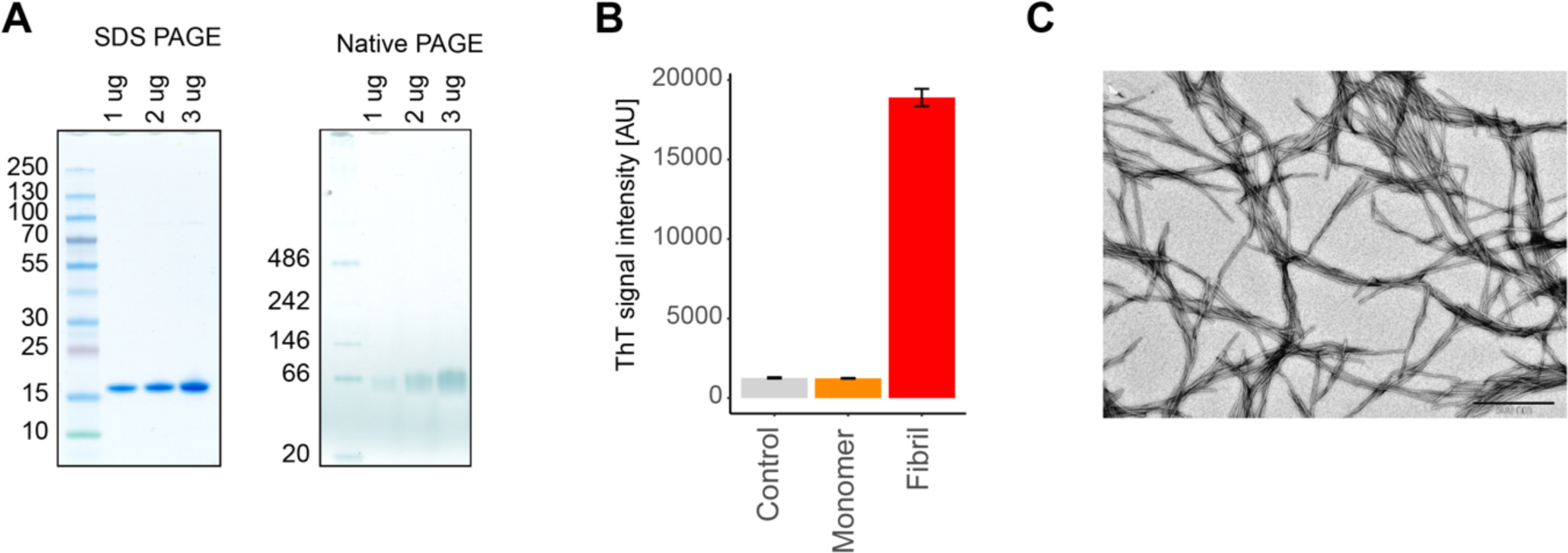
Quality controls α-Synuclein monomer and fibrils. **A** SDS PAGE and Native PAGE of monomeric α-Synuclein. **B** ThT intensity of Control, monomeric-and fibrillary α-Synuclein. **C** TEM image of the α-Synuclein fibrils used.

**Supplementary figure 8.**
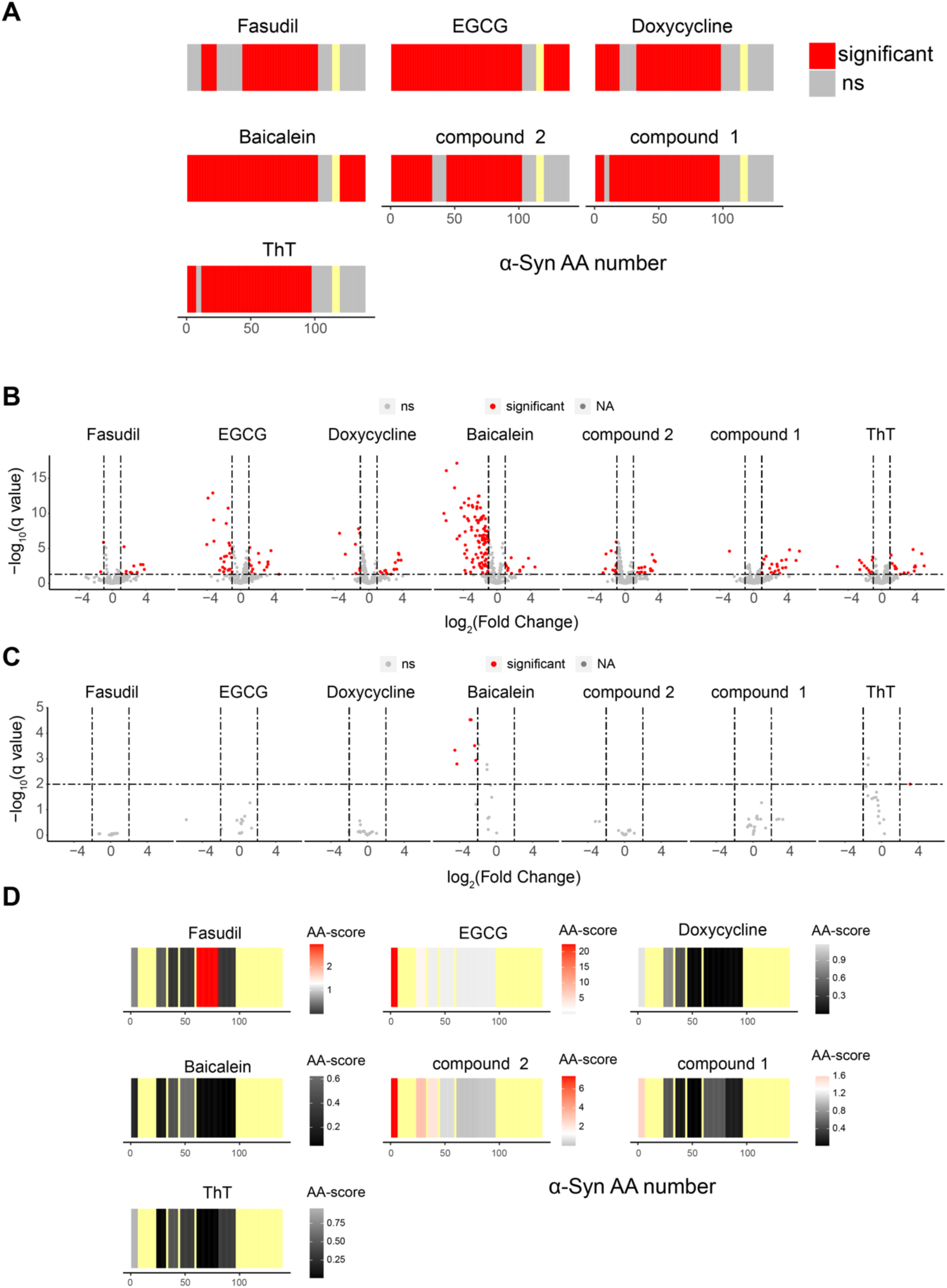
Compound interactions with α-Synuclein monomer. **A, B** LiP peptide fingerprints (non-significant in light grey, significant in red, not detected in yellow) (**A**) and volcano plots of LiP peptide intensities (significant in red, non-significant in grey) **(B**) for α-Synuclein monomer treated with compounds compared to treatment with the DMSO control. **C** Volcano plot comparing the control (i.e., trypsin-only) peptide intensities of compound-treated α-Synuclein monomer to DMSO-treated monomer; colours as in B. **D** LiP peptide fingerprints of α-Synuclein treated with compounds and normalized for the trypsin-only control data (The scale indicates the score per peptide. The significance threshold of −log_10_(0.05) x log_2_(2) is shown in white, with red indicating higher scores. The more intense the red colour, the higher the score. Not significant in grey. Not detected in yellow).

**Supplementary figure 9.**
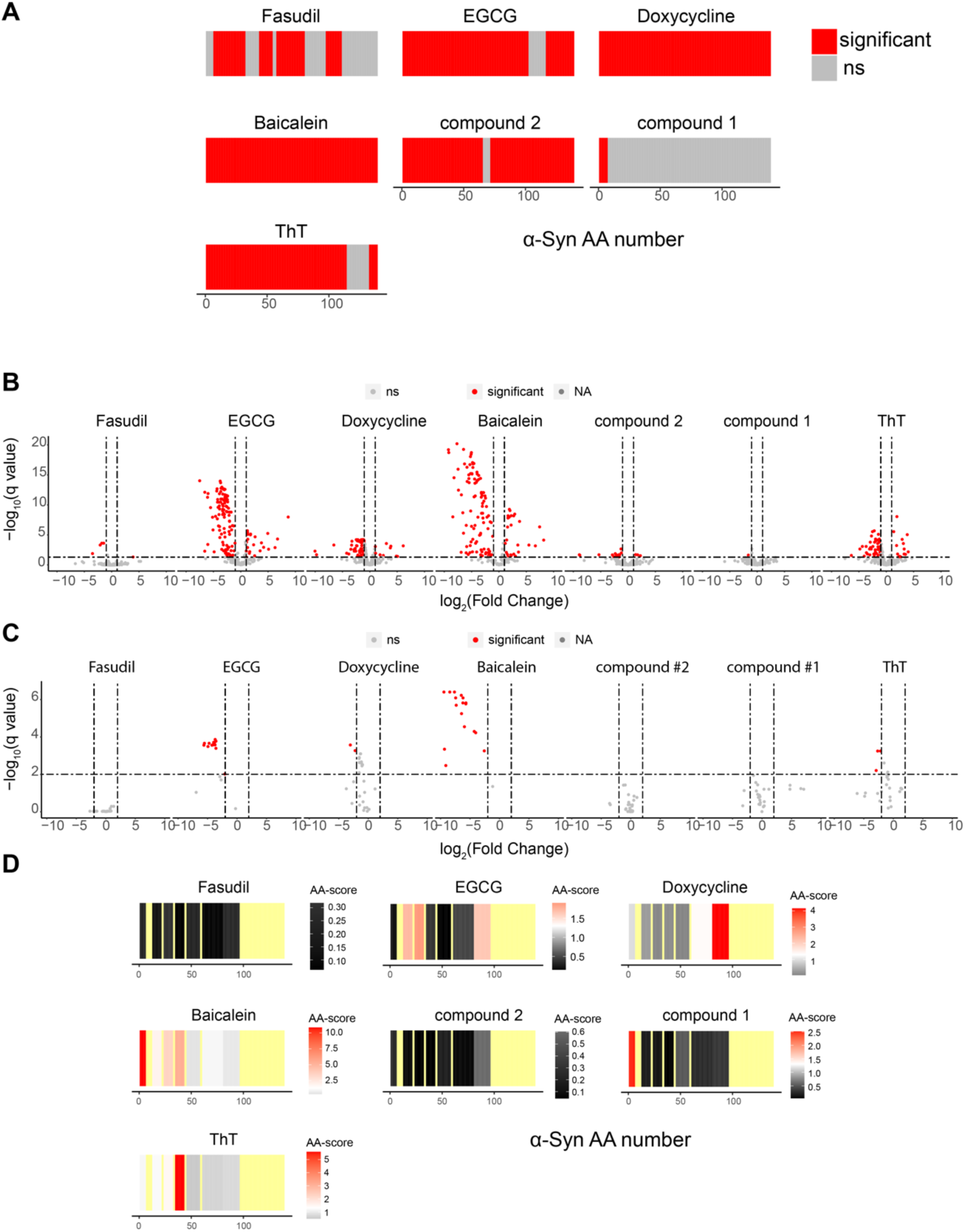
Compound interactions with α-Synuclein fibril. **A** Peptide analysis fingerprints of α-Synuclein treated with compounds (non-significant in light grey, significant in red, not detected in yellow). **B** Volcano plot of compound treated α-Synuclein fibril (non-significant in light grey, significant in red). **C** Volcano plot of the control intensities of compound treated α- Synuclein fibril (non-significant in light grey, significant in red). **D** Peptide fingerprint of the tryptic control normalised data (The scale indicates the score per peptide. The significance threshold of −log_10_(0.05) x log_2_(2) is shown in white, with red indicating higher scores. The more intense the red colour, the higher the score. Not significant in grey. Not detected in yellow).

**Supplementary figure 10.**
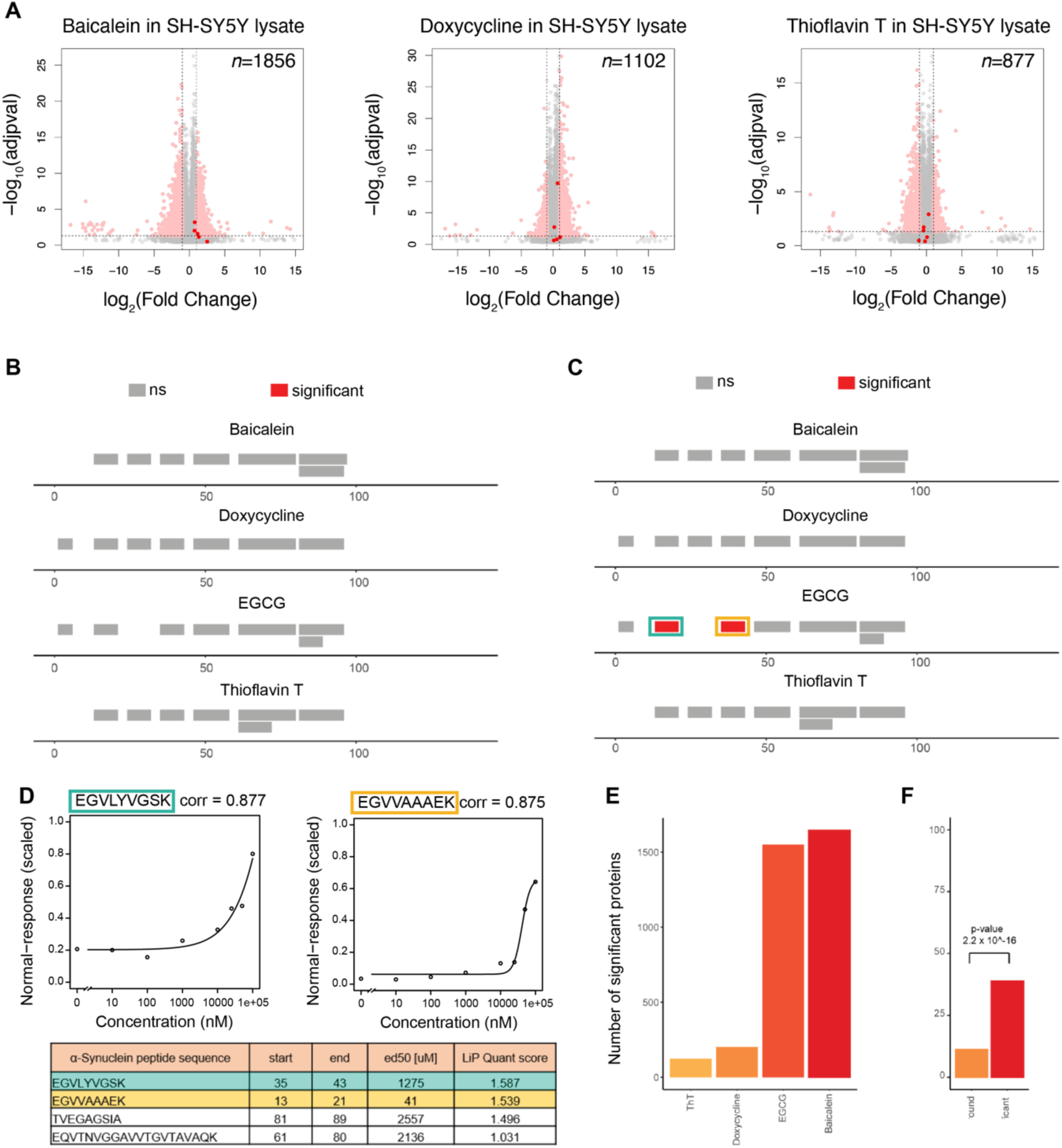
**A.** Volcano plots showing peptides with altered abundance after spike-in of the indicated compounds into cell lysates. Number of significant hits and total number of detected peptides are indicated. **B, C** The plots show a LiPQuant analysis of alpha-synuclein in the presence of indicated compounds, with detected peptides colored according to the LiPQuant score. Red is significant. Grey is not significant. Note that A and B are scaled differently; LiPQuant scores are shown to a threshold of 2.0 (B) and 1.5 (C). **D** The table shows α-Synuclein peptides and their start and end position, ec50 value and LiPQuant score. The blue and yellow colors indicate the two α- Synuclein peptides with LiPQuant score > 1.5 in the presence of EGCG. The plots show curve fits of peptide intensity versus EGCG concentration for the two α-Synuclein peptides with LiPQuant score > 1.5 in the presence of EGCG. **E.** Number of significant proteins in the presence of the indicated compounds, at LiPQuant score > 1.5. **F**. Fisher’s exact test of significant proteins (LiPQuant score > 1.5) detected in the presence of Doxycycline and in the Doxycycline pulldown experiment compared to not significant background.

**Supplementary figure 11.**
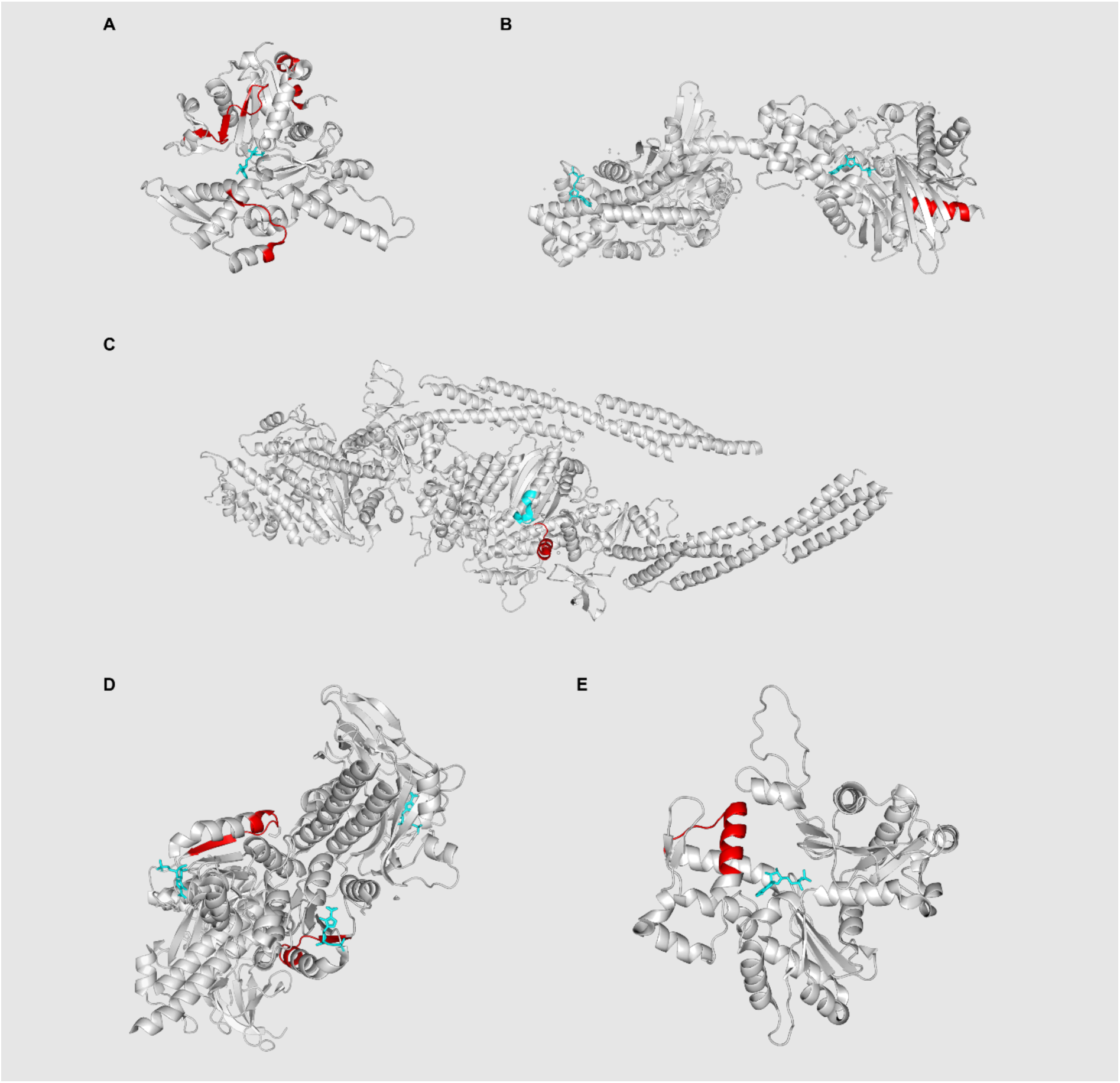
Significant hits with ATP/ADP/C2R bound pdb structures. **A** ACTR (pdb: 6uhc). Significant hits in red, ATP in cyan. **B** HK1 (pdb: 1dgk). Significant hit in red, ADP in blue. **C** MYH10 (pdb: 4pd3). Significant hit in red. ATP binding site from “uniprot.org” in cyan. **D** PAICS (pdb: 6yb8). Significant hit in red, C2R in cyan. **E** ACTB (pdb: 3j82). Significant hit in red, ADP in cyan.

**Supplementary figure 12.**
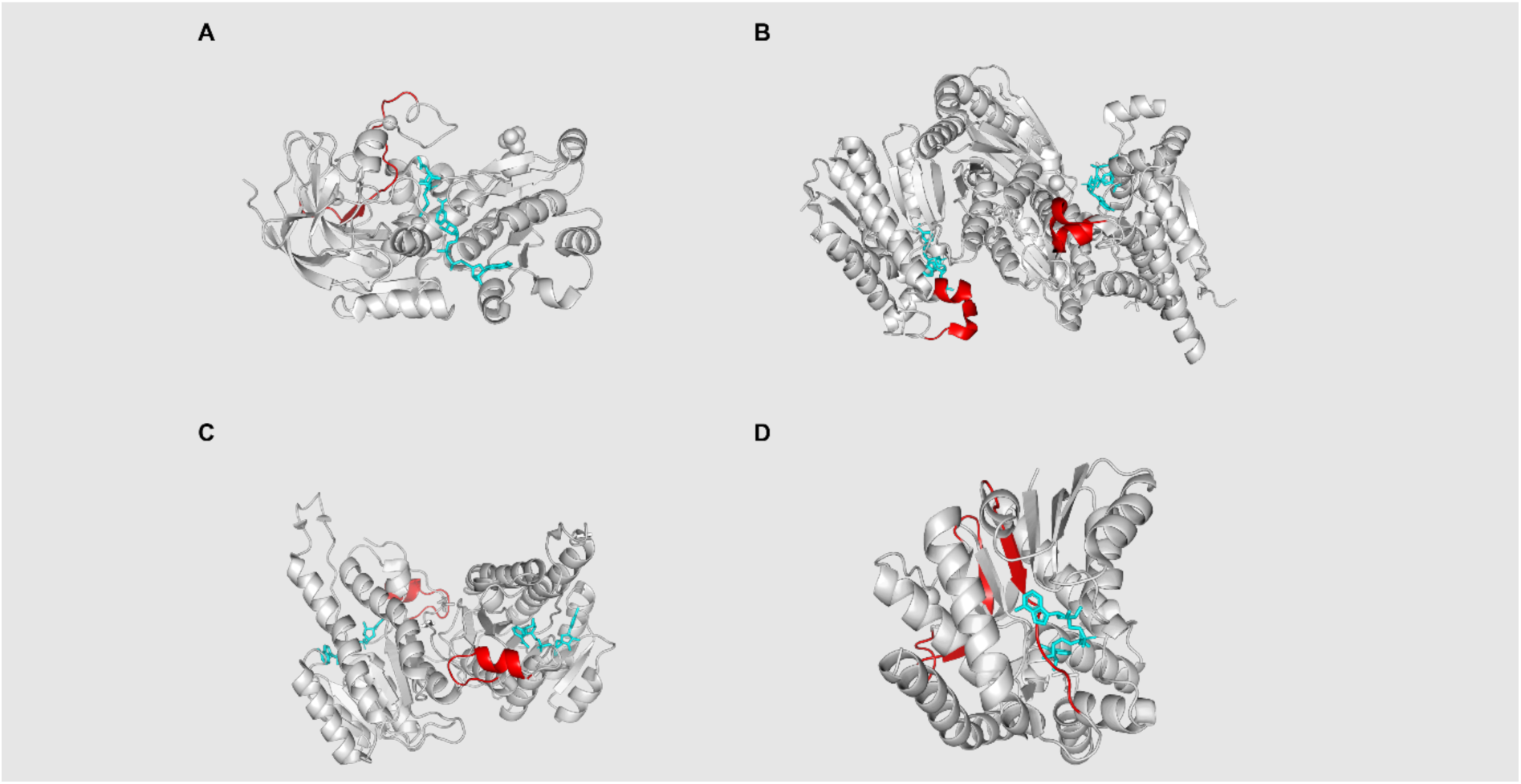
Significant hits with NAD/NDP bound pdb structures. **A** ADH5 (pdb: 1mc5). Significant hit in red, NAD and AHE in cyan. **B** IDH2 (pdb: 4ja8). Significant hit in red, NDP in cyan. **C** HSD17B10 (pdb: 2o23). Significant hit in red, NAD in cyan. **D** MDH2 (pdb: 2dfd). Significant hit in red, NAD in cyan.

